# Bacterial evolution in high osmolarity environments

**DOI:** 10.1101/2020.05.07.081752

**Authors:** Spencer Cesar, Maya Anjur-Dietrich, Brian Yu, Ethan Li, Enrique Rojas, Norma Neff, Tim F. Cooper, Kerwyn Casey Huang

## Abstract

Bacteria must maintain a cytosolic osmolarity higher than that of their environment in order to take up water. High osmolarity environments therefore present a formidable stress to bacteria. To explore the evolutionary mechanisms by which bacteria adapt to high osmolarity environments, we selected *Escherichia coli* in media with a variety of osmolytes and concentrations for 250 generations. Adaptation was osmolyte-dependent, with sorbitol stress generally resulting in increased fitness in conditions with higher osmolarity, while selection in high concentrations of proline resulted in increased fitness specifically on proline. Consistent with these phenotypes, sequencing of the evolved populations showed that passaging in proline resulted in specific mutations in an associated metabolic pathway that increases the ability to utilize proline for growth, while evolution in sorbitol resulted in mutations in many different genes that generally improve growth in high osmolarity conditions at the expense of growth at low osmolarity. High osmolarity decreased growth rate but increased mean cell volume compared with growth on proline as the sole carbon source, demonstrating that osmolarity-induced changes in growth rate and cell size follow an orthogonal relationship from the classical Growth Law relating cell size and nutrient quality. Isolates from a sorbitol-evolved population that capture the likely temporal sequence of mutations revealed by metagenomic sequencing demonstrate a tradeoff between growth at high and low osmolarity. Our study highlights the utility of experimental evolution for dissecting complex cellular networks and environmental interactions, particularly in the case of behaviors that can involve both specific and general metabolic stressors.

**Importance:** For bacteria, maintaining higher internal solute concentrations than the environment allows cells to take up water. As a result, survival is challenging in high osmolarity environments. To investigate how bacteria adapt to high osmolarity environments, we evolved *Escherichia coli* in a variety of high osmolarity solutions for hundreds of generations. We found that evolved populations adopted different strategies to improve their growth depending on the osmotic passaging condition, either generally adapting to high osmolarity conditions or better metabolizing the osmolyte as carbon source. Single-cell imaging demonstrated that enhanced fitness was coupled to faster growth, and metagenomic sequencing revealed mutations that reflect growth tradeoffs across osmolarities. Our study demonstrates the utility of long-term evolution experiments for probing adaptation during environmental stress.

## Introduction

Osmolarity is a fundamental and dynamic physiochemical property of microbial environments. Bacteria inhabit a wide range of osmotic niches that range from very dilute environments (e.g. fresh water) to highly concentrated environments (e.g. soil, salt marshes). Halophiles live in highly saline environments (above ~20% salinity), where many exhibit slow growth relative to enteric bacteria such as *Escherichia coli* (1, 2). Bacteria must also cope with large fluctuations in their osmotic environment that occur over a range of timescales. For example, enteric bacteria often transition between the high osmolarity intestinal environment and low-osmolarity environments outside the host, potentially experiencing an acute hypoosmotic shock upon exit from the host (3). Within the small intestine itself, luminal osmolarity undergoes large variations during digestion as macromolecules are metabolized and crypt cells actively secrete electrolytes that contribute to luminal water secretion (4). Bacteria in the large intestine experience frequent osmotic shifts, as epithelial cells actively transport sodium ions into the digestive cavity to aid in water absorption (5). Thus, the molecular mechanisms by which bacteria adapt to the extracellular osmotic properties are a key determinant of their ability to establish and maintain populations across a range of environments.

Bacteria maintain a large (~1 atm) cytosolic osmotic pressure relative to their environment that necessitates a rigid cell envelope including a cell wall to prevent lysis due to the resulting turgor pressure (6, 7). Most (if not all) bacterial species possess several pathways devoted to osmoregulation and use compatible solutes such as proline and glycine betaine as major regulators of osmotic pressure (8–10). The steady-state growth rate of bacterial cells is approximately inversely correlated with medium osmolarity (11). This relationship does not depend on the osmolyte used to modulate osmolarity, indicating that the growth rate decrease is not a result of specific toxicity (12, 13). Hyperosmotic stress impacts growth rate by decreasing the ribosomal elongation rate during translation (14), and could also inhibit specific metabolic pathways and/or decrease the partial vapor pressure of water within the cytoplasm, thereby having detrimental consequences on the efficiency of biochemical reactions(15). The global nature of osmotic stress makes it likely that a variety of mutations can confer comparable fitness advantages and thus it is difficult to pinpoint specific genes that impact fitness in high osmolarity environments, making experimental evolution an attractive methodology for probing adaptive responses.

In the Gram-negative, rod-shaped enteric bacterium *Escherichia coli*, turgor pressure is estimated to be ~0.3-1 atm and may depend on the growth medium (16, 17). Longitudinal stretching of the cell wall decreases with increasing osmolarity, although there remains a large elastic stretching (indicating positive turgor pressure) at osmolarities that are prohibitive to growth (18). The rate of cell-wall expansion in *E. coli* is not directly dependent on turgor pressure; hyperosmotic shock does not affect the instantaneous growth rate of single cells on time scales of minutes, indicating that cells can buffer against osmotic changes to continue fast growth (18). This buffering capacity suggests that cells could evolve to grow at a faster rate in higher osmolarity environments.

The ability to conduct rigorous evolution experiments in laboratory environments has been exploited across a wide variety of organisms, including plants, vertebrates, and microorganisms (19). Microorganisms are uniquely suited for long-term evolution studies due to the ease with which they can be propagated, stored, and genetically manipulated (19–21). Experimental evolution provides a means for revealing the capacity for improved growth at high osmolarity, and the dependence of growth behaviors on the specific osmolyte. Moreover, the genetic basis for changes in growth during evolution in high osmolarity can elucidate the mechanisms by which cells mitigate the broad detrimental effects of high intracellular osmolarity.

In this study, we selected *E. coli* populations for hundreds of generations in high osmolarity media supplemented with various osmolytes, and identified phenotypic adaptations of these evolved populations across a range of osmolyte concentrations. We found that the evolved populations possessed a wide range of fitness advantages that were osmolyte-specific. In particular, we found that proline stress primarily resulted in increased fitness when grown on proline as a carbon source, while sorbitol stress generally resulted in increased fitness across a wide range of concentrations of multiple osmolytes. Passaging in the other osmolytes did not result in any adaptation over the same number of generations. Sequencing of the evolved populations revealed mutations that broadly conferred adaptive benefits to different osmotic environments as well as tradeoffs, and that altered growth rate and cell size in osmolyte-dependent manners.

## Results

### Osmolyte-dependent phenotypic changes over hundreds of generations of passaging

To ascertain whether and how *E. coli* can adapt to growth in high osmolarity environments, we propagated populations in DM25 (Davis Minimal + 25 mg/L glucose) medium (Methods) using four replicates with each of five common osmolytes: glycine betaine, proline, sorbitol, sucrose, and sodium chloride (Fig. 1A). To avoid adaptation specific to growth in DM25, we selected as the ancestral strain TC1407, a derivative of *E. coli* REL606, which had already been propagated in glucose-supplemented medium for 4,000 generations and therefore had already adapted to growth in this minimal medium. Glycine betaine and proline were chosen as osmolytes based on their well-known role in *E. coli* osmoregulation as osmoprotectants, which could impact which mutations are adaptive (8, 9). Sucrose and sorbitol are sugars that *E. coli* does not preferentially metabolize, although sorbitol can be metabolized by certain laboratory strains, including TC1407 (22). Both of these sugars have been previously studied in the context of growth at high osmolarity and result in specific changes in *E. coli* gene expression (23, 24). Sodium chloride is a salt that has been used previously to apply osmotic shock to cells and can also impact other cellular functions by improving acid tolerance, altering lipid bilayer balance, and altering gene expression (25–27). Two concentrations for each osmolyte were chosen for passaging to probe a range of behaviors of the ancestor (Fig. 1B-F). As both sorbitol (Fig. 1C) and proline (Fig. 1F) can be used as a carbon source by TC1407 cells, these conditions supported more growth than in glucose alone. Nonetheless, growth rate was slower at higher osmolarity for all osmolytes (Fig. 1B-F), as expected based on previous studies (11). Cell densities were somewhat lower at higher concentrations of glycine betaine (Fig. 1B) and sodium chloride (Fig. 1D) compared to growth without the osmolyte, suggesting that the stress posed by high concentrations of these osmolytes impairs efficient carbon utilization. As a control, we propagated replicate populations in DM25 alone. All populations were evolved in their selective environment for a total of 250 generations (38 daily 1:100 dilution transfer cycles). To simplify presentation and discussion of our results, we will refer to population *i* (=1, 2, 3, 4) evolved in medium M with concentration *x* after 250 generations as M*x*−*i*, where M=Pr for proline, So for sorbitol, Gl for glycine betaine, Na for sodium chloride, and Su for sucrose.

**Figure 1:**
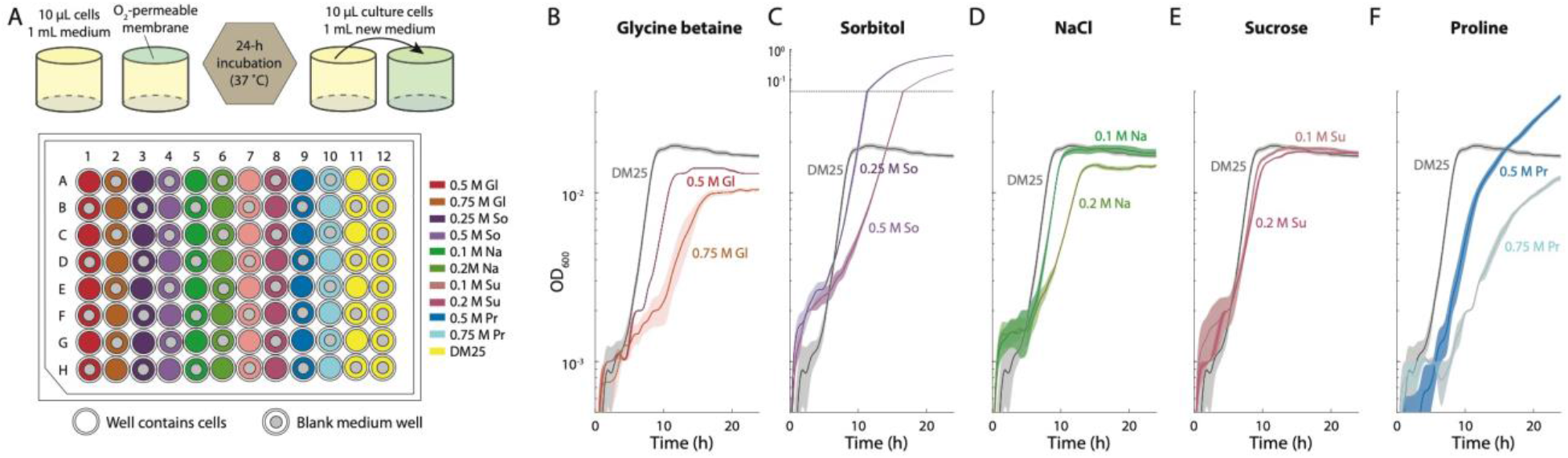
Schematic of evolution experiment across passaging conditions that result in a wide range of ancestral growth behaviors. A) The ancestral strain (TC1407) was passaged daily with 1:100 dilutions as four independent populations in each of 11 environments of DM25 medium supplemented with osmolytes as denoted on the plate. Empty wells were used as negative controls. Gl: glycine betaine; Na, sodium chloride; Pr, proline; So, sorbitol; Su, sucrose. B-F) Higher osmolarity generally inhibits growth, although sorbitol and proline can be utilized as carbon sources. TC1407 cells were grown in the passaging concentrations of glycine betaine (B), sorbitol (C), NaCl (D), sucrose (E), and proline (F). For all osmolytes, growth during the first 10 h was inhibited more at higher osmolyte concentrations. Glycine betaine and NaCl also decreased the carrying capacity, signifying less efficient use of glucose. The carrying capacity was greatly increased in sorbitol and in 0.5 M proline due to utilization of the osmolyte as a carbon source. Growth in sucrose was relatively unaffected by osmolarity. Growth curves are averages of *n*=4 replicates, and shaded regions represent the standard error of the mean (SEM). The SEM is small at some time points and is partially obscured by the lines.

To quantify adaptation in the evolved populations, we measured growth curves after 250 generations of passaging and compared to those of the ancestor in each environment (Fig. 2A, B, S1). For consistency, populations were revived from a −80 °C frozen stock and equilibrated in DM25 via three 24-h passages before growth-curve measurements. These passages were critical for the reproducibility of measurements, as growth from the frozen stock did not equilibrate until the third passage. We did not perform a passage in the test environment prior to growth-curve measurements so that cells were in the same physiological state at the start of each fitness measurement. Passage in DM25 alone did not result in noticeable changes in growth in any of the evolved populations (Fig. S1A), suggesting that 250 generations was not sufficient for measurable adaptation in the base environment lacking any added osmolyte. Any changes in growth parameters selected in the presence of an osmolyte were thus likely to be due specifically to the difference in medium.

**Figure 2:**
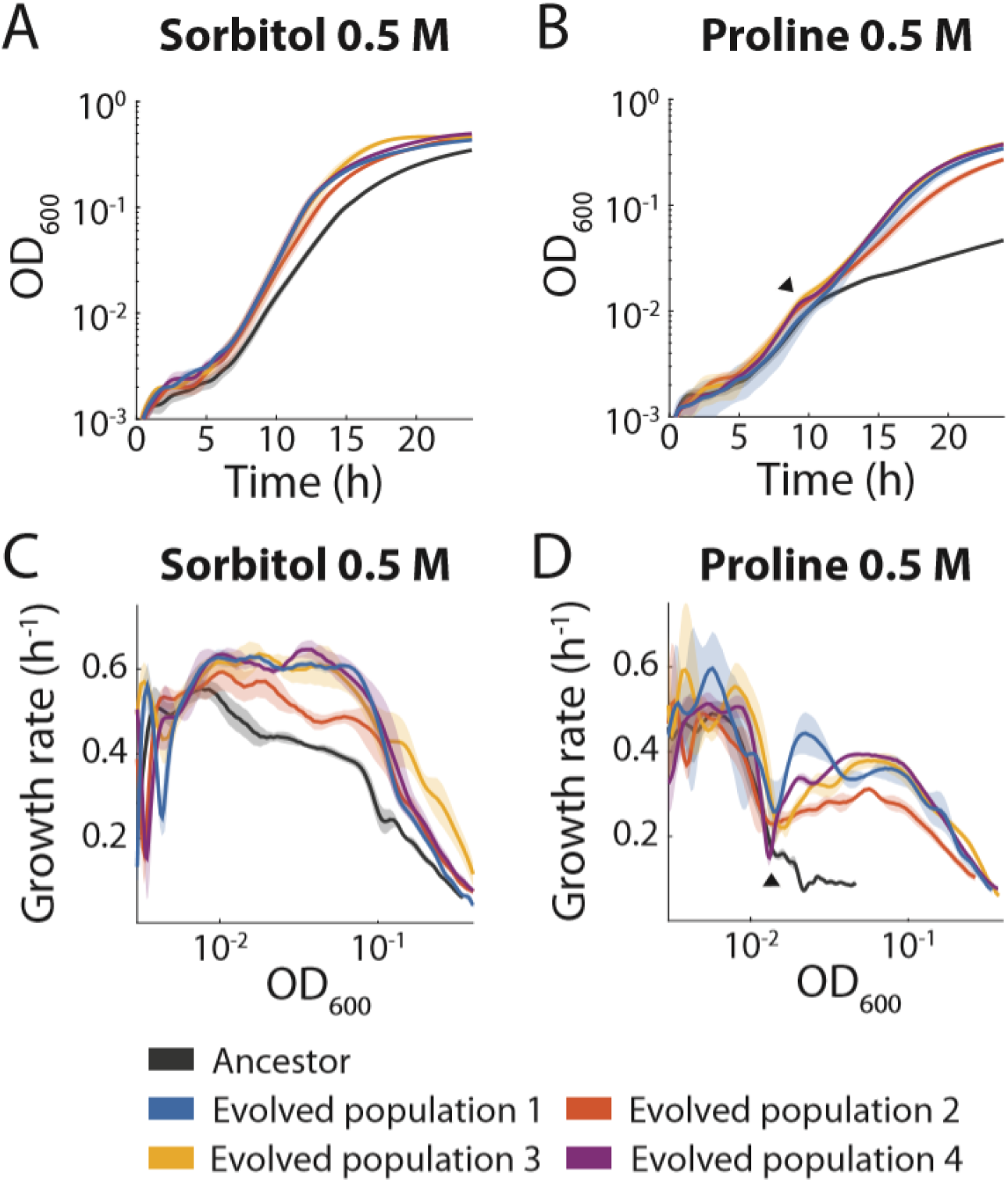
All populations evolved in 0.5 M proline or sorbitol grow better than the ancestor in their passaging medium. A,B) The populations evolved in 0.5 sorbitol (A) or 0.5 M proline (B) grew faster and to a higher OD after 24 h than the ancestor in their passaging medium. For both osmolytes, there was a range of growth behaviors of the four populations, suggesting different adaptations. Growth curves are averages of *n*=3 replicates for each of the evolved populations and averages of *n*=6 replicates for the ancestor, with shaded regions representing the standard error of the mean (SEM). The black arrowhead in (B) indicates when the cultures likely shift from glucose to proline utilization. The SEM is small at some time points and is partially obscured by the lines. C,D) To account for the different growth kinetics of the ancestor and evolved populations in (A,B), we analyzed growth rate as a function of OD_600_. For sorbitol (C) and proline (D), growth rate was consistently higher in the evolved populations. The black arrowhead marks the same time as in (B), when growth rate briefly decreases. The SEM is small at some time points and is partially obscured by the lines.

While there was little to no change in growth in the populations passaged in 0.25 M sorbitol (Fig. S1D), the four populations passaged in 0.5 M sorbitol had substantially higher growth rates (0.52-0.65 h^−1^ at OD=0.2) in DM25 + 0.5 M sorbitol compared with the ancestor (0.43±0.01 h^−1^ at OD=0.2, errors are standard error of the mean (SEM) unless otherwise noted; Fig. 2A, C). A large fitness advantage over the ancestor was apparent in the evolved populations from both proline concentrations (Fig. 2B, D, S1H), due to faster growth rate after an apparent shift from glucose to proline consumption at ~10 h or ~15 h in the 0.5 M or 0.75 M condition, respectively. After 10 h, the populations passaged in 0.5 M proline had maximum growth rates in DM25 + 0.5 M proline between 0.31 h^−1^ and 0.44 h^−1^, while the ancestor grew at ~0.10±0.01 h^−1^.

Because the growth dynamics were quite different between the evolved populations and the ancestor across the growth curve, we examined the relationship between instantaneous growth rate and OD, which are generally closely related across mutants with the same maximal growth rate even when the lag phase varies (28). The evolved populations in 0.5 M sorbitol and 0.5 M proline consistently exhibited higher growth rates in their passaging medium than the ancestor at similar ODs (Fig. 2C, D). There were little to no changes in growth following selection in glycine betaine (Fig. S1B, C), NaCl (Fig. S1E), or sucrose (Fig. S1F, G) environments; the concentration and/or stressful impact of the osmolyte or the lower cell density relative to proline and sorbitol culturing may explain the lack of adaptation in these conditions. Nonetheless, proline- and sorbitol-mediated adaptation demonstrates that *E. coli* can increase its growth rate in high osmolarity environments.

### Evolution in sorbitol led to general fitness increases in high osmolarity environments

One 0.5 M sorbitol-evolved and one 0.5 M proline-evolved population were chosen for further assessment to determine the nature of the observed changes in fitness (hereafter called So0.5-1 and Pr0.5-1). To probe whether the increased fitness of the So0.5-1 population relative to the ancestor in DM25+0.5 M sorbitol extended to other osmolarities, we examined growth in DM25 media with a range of concentrations of sorbitol. We first passaged each population three times in DM25 for 24 h, and then measured growth curves for 72 h. We defined the relative fitness between the evolved population and the ancestor as the ratio of the ODs of the evolved population and the ancestor when the evolved population reached half its final OD (either at saturation or the last OD measurement if the culture did not reach saturation) (Fig. 3A). The rationale for this metric is that some of the cultures at high osmolarity took >72 h to saturate, at which point evaporation prevented accurate growth measurements; in all cases, visual inspection of the growth curves led to conclusions in agreement with our relative fitness calculations.

**Figure 3:**
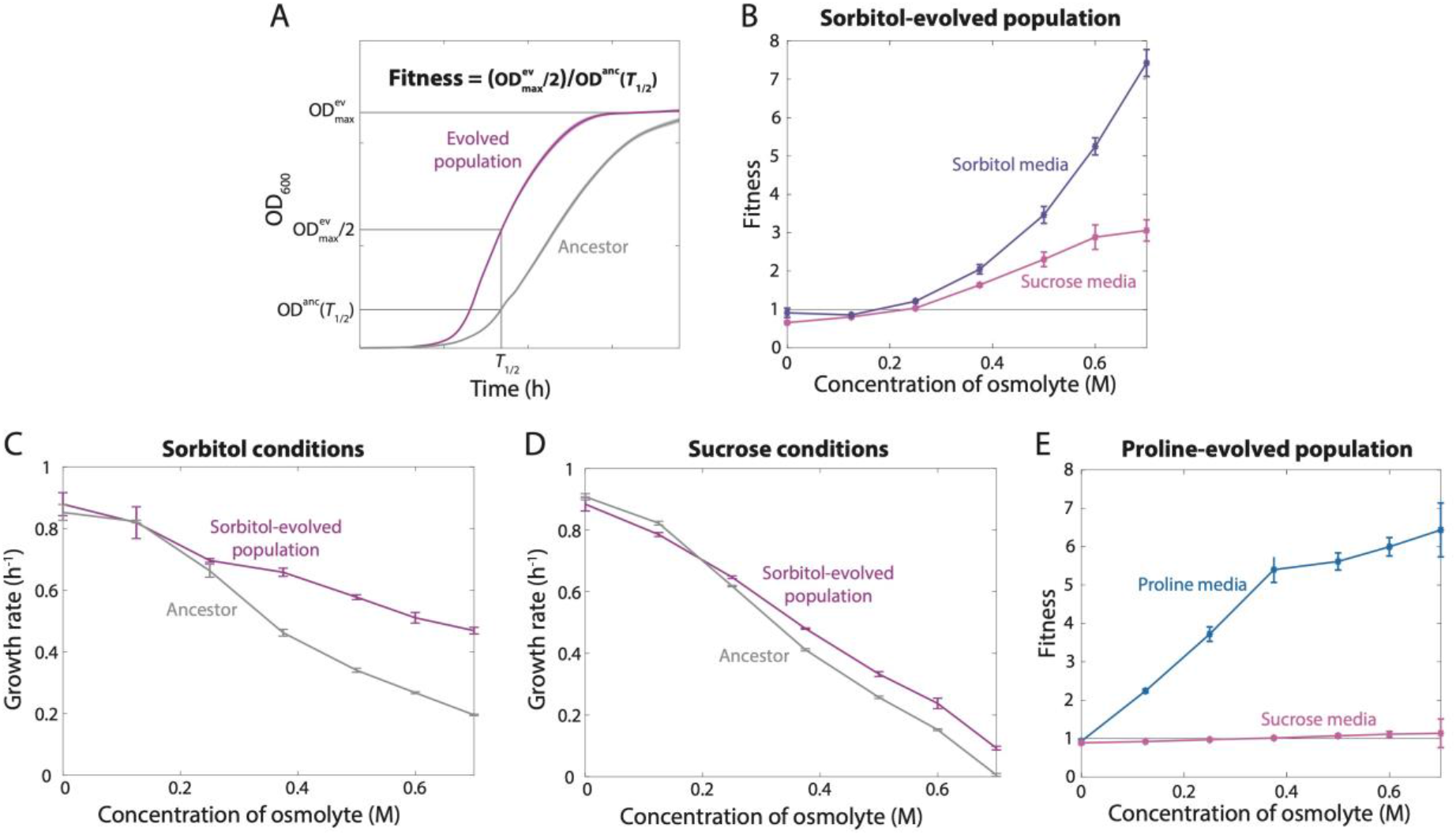
Sorbitol-evolved populations have increased fitness at high osmolarity, while proline-evolved populations have increased fitness specifically in proline. A) Schematic of calculation of relative fitness from growth curves (Fig. S2), which we define as the ratio of half the saturation OD of an evolved population 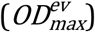 to the OD of the ancestor (OD^anc^) at the same time point (*T*_1/2_). B) The So0.5-1 population had higher and monotonically increasing fitness relative to the ancestor in both sorbitol (purple) and sucrose (pink) for concentrations ≥0.25 M, and slightly reduced fitness in DM25 or DM25+0.125 M osmolyte, indicating that the population has adapted to higher osmolarity conditions (*n* = 4). C,D) The maximum growth rate of the ancestor and the sorbitol-evolved populations in sorbitol (C) and sucrose (D) decreased steadily with increasing osmolarity, with a faster decrease in the ancestor leading to larger growth rate differences at higher osmolarities (*n* = 4). E) The Pr0.5-1 population exhibited monotonically increasing fitness relative to the ancestor at all concentrations of proline (blue), and neutral fitness without proline. Growth in sucrose (pink) was virtually identical to the ancestor at all concentrations (*n* = 4).

The So0.5-1 population had a clear relative fitness advantage over the ancestor at sorbitol concentrations ≥0.25 M, with the advantage increasing with concentration, even above 0.5 M (Fig. 3B). Interestingly, the So0.5-1 population maintained a density of OD_600_ >0.45 across all conditions with added sorbitol, while the density of the ancestor decreased dramatically in media with 0.7 M sorbitol (Fig. S2A). Without added sorbitol (DM25), the ancestor and the evolved population had maximum growth rates within 4% of each other (Fig. 3C). In DM25+0.125 M sorbitol, the evolved population and ancestor also had similar maximum growth rates (0.82 ±0.05 h^−1^ for the evolved population vs. 0.82±0.01 h^−1^ for the ancestor); however, after a peak in growth rate early in the passage cycle, the evolved population exhibited a consistently lower growth rate than the ancestor (Fig. S2A), consistent with the small relative fitness decrease compared with the ancestor at this sorbitol concentration (Fig. 3B). With ≥0.25 M added sorbitol, the So0.5-1 population had a higher maximum growth rate than the ancestor, with the difference increasing with increasing sorbitol concentration due to the more rapid decrease in growth rate of the ancestor (Fig. 3C). These conclusions were confirmed by fitting the growth curves to a Gompertz function (Fig. S3). Thus, the enhanced relative fitness of the So0.5-1 population at osmolarities ≥0.25 M is likely due to insulation from the detrimental effects of high osmolarity experienced by the ancestor.

To test whether the relative fitness advantages during growth with sorbitol were also present at high concentrations of other osmolytes, as opposed to specific adaptations involving sorbitol protection and/or utilization, we examined growth of the So0.5-1 population in a range of sucrose concentrations. Sucrose was chosen to assess nonspecific adaptation since most *E. coli*, including our ancestor, do not metabolize sucrose and do not use sucrose for osmoregulation. After multiple passages in DM25, cells were grown in media with sucrose added to DM2500 (DM+2500 mg/L glucose), which supports higher carrying capacities than DM25 (Fig. S2A, B) to more closely match those of growth in DM25+sorbitol (Fig. S2A) and hence better facilitate detection of fitness differences. In sucrose, the So0.5-1 population displayed a similar pattern as in sorbitol, with a monotonically increasing relative fitness advantage compared with the ancestor at 0.25 M and above (Fig. 3B). The fitness advantage manifested early in the growth curve before glucose was exhausted in both sorbitol and sucrose (Fig. S2A, B), providing further support that the adaptations are not specific to sorbitol as a carbon source. There was again a small decrease in relative fitness below 0.25 M sucrose, indicating that the population had adapted to higher osmolarities at the expense of growth in low osmolarity (Fig. 3B). As in sorbitol, the maximum growth rate of the evolved population was greater than that of the ancestor for all concentrations ≥0.25 M sucrose and the difference increased with concentration, but the ancestor grew faster in 0.125 M sucrose (Fig. 3D) (*p*=0.01, 2-tailed t-test). The evolved population also had growth advantages over the ancestor in DM2500 supplemented with high concentrations of proline, glycine betaine, and NaCl (Fig. S4A-E), indicating a general fitness benefit at high osmolarity. The evolved population also had increased fitness relative to the ancestor in DM2500 with 0.5 M sorbitol (Fig. S4F), a condition with excess glucose such that metabolism of sorbitol is not necessary for growth to high density. Finally, we tested growth in DM25 supplemented with 0.4 M sorbitol or sucrose and 0.1 M proline, to determine if the fitness advantage depended on the absence of an osmoprotectant (Fig. S5A, B). While growth of both the ancestor and the evolved population was faster with addition of proline, the evolved population still exhibited an advantage over the ancestor. Thus, the So0.5-1 population generally grows faster in media supplemented with sufficiently concentrated osmolytes, with a tradeoff at lower osmolarities.

### Evolution in proline led to proline-specific fitness increases at all osmolarities

We next examined growth of the Pr0.5-1 population, which grew to a higher OD after 24 h than the ancestor across a range of proline concentrations (Fig. S2C). In DM25 without added osmolyte, the Pr0.5-1 population grew very similarly to the ancestor, indicating that adaptation to growth on proline did not exert a fitness cost during growth on glucose. By contrast to the So0.5-1 population grown in sorbitol, the early stages of growth of the Pr0.5-1 population in DM25+proline were quantitatively similar to those of the ancestor until a growth shift at ~8-12 h that is likely due to transitioning from glucose to proline utilization (Fig. S2C); it was only after the shift that the evolved population exhibited clear advantages (Fig. S2C). Using the same fitness metric (Fig. 3A) as for sorbitol, the Pr0.5-1 population had an advantage over the ancestor that increased monotonically with increasing proline concentration (Fig. 3E). Notably, the evolved population achieved higher OD values after 24 h than the ancestor on all proline concentrations >0.125 M, suggesting that it utilizes proline as a carbon source more efficiently (Fig. S2C). The carrying capacity of the evolved population decreased more slowly above 0.25 M than the ancestor (Fig. S2C).

After the shift to proline utilization in 0.125 M proline, the maximum growth rate of the evolved population was significantly higher than that of the ancestor (0.50±0.004 h^−1^ vs. 0.31±0.003 h^−1^; *p*<10^−4^, two-tailed t-test); in 0.25 M proline, the difference in maximum growth rates was increased (0.50±0.002 h^−1^ vs. 0.22±0.001 h^−1^; *p*<10^−4^, two-tailed t-test). At higher concentrations of proline, the maximum growth rates of the evolved population and the ancestor both decreased, with the ancestor experiencing a larger fractional decrease in its growth rate with increasing concentration compared with the evolved population (Fig. S2C), leading to increased relative fitness (Fig. 3E).

To resolve whether the increased growth of the evolved population at high proline concentrations was due to enhanced ability to deal with osmotic stress, as opposed to a metabolic advantage, we grew the Pr0.5-1 population in a range of sucrose concentrations. By contrast to the So0.5-1 population grown in sucrose, the fitness (Fig. 3E) and the growth rate (Fig. S2D) of the Pr0.5-1 population closely mirrored that of the ancestor at all concentrations. Taken together, these data indicate that the Pr0.5-1 population adapted specifically to growth on proline rather than to high osmolarity.

### Sorbitol- and proline-evolved populations exhibit osmolyte-dependent changes in steady-state growth rate

During a passage cycle in DM25+0.5 M sorbitol or proline, cells undergo complex growth dynamics involving exit and re-entry into stationary phase, as well as a shift from early utilization of glucose to growth on sorbitol or proline. Thus, we sought to determine whether the increases in fitness in high osmolarity conditions could be attributed specifically to increased growth rate, as opposed to a more complex set of drivers associated with the passaging. We measured single-cell growth rates during steady-state, exponential growth (Methods) in DM2500, DM2500+0.5 M sucrose, and DM (no glucose) +0.125 M proline. In the first two cases, DM2500 was used as a culture medium because the low density of DM25 cultures makes it difficult to acquire data for a large population of cells. In DM2500, Pr0.5-1 cells exhibited a unimodal distribution of growth rates quantitatively matching that of the ancestor (*p*=0.61, two-tailed t-test) (Fig. 4A), while the mean growth rate of So0.5-1 cells was significantly lower than the ancestor by 12% (*p*=8.1×10^−4^, two-tailed t-test) (Fig. 4A). These observations are consistent with our population growth measurements, in which the Pr0.5-1 population grew similarly to the ancestor in DM2500 (Fig. 3E, S2D) while the So0.5-1 population had a small disadvantage (Fig. 3B, S2B).

**Figure 4:**
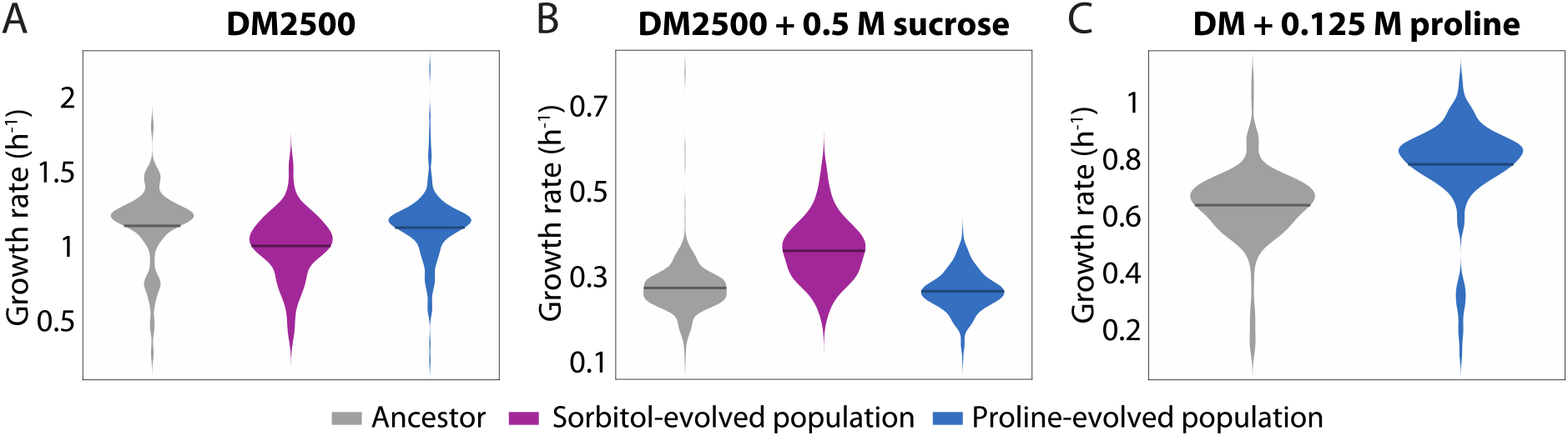
Fitness advantages relative to the ancestor are correlated with increases in steady-state growth rates of the sorbitol- and proline-evolved populations. A) Cells from the So0.5-1 population grew slower than the ancestor in DM2500 at steady-state (*p*=8.1×10^−4^, *t*-test) while Pr0.5-1 cells exhibited similar growth rates as the ancestor (*p*=0.61, *t*-test), consistent with fitness measurements in Fig. 3B and 3E, respectively. The growth rate for each cell was defined as its mean instantaneous growth rate over 15 min of imaging (*n*>77 cells for the ancestor, *n*>97 cells for the sorbitol-evolved population, and *n*>218 cells for the proline-evolved population). Horizontal lines represent means across cells. B) So0.5-1 cells grew more quickly at steady state in DM2500+0.5 M sucrose than the ancestor (*p*<10^−4^, *t*-test) or Pr0.5-1 cells (*p*<10^−4^, *t*-test), which had similar growth rates, again consistent with fitness measurements. The growth rate for each cell was defined as its mean instantaneous growth rate over 21 min of imaging (*n*>405 cells for the ancestor, *n*>125 cells for the sorbitol-evolved population, and *n*>291 cells for the proline-evolved population). Horizontal lines represent means. C) Pr0.5-1 cells grew more quickly at steady state in DM+0.125 M proline than the ancestor (*p*<10^−4^, *t*-test), demonstrating that passaging in proline led to enhanced ability to metabolize proline. The growth rate for each cell was defined as its mean instantaneous growth rate over 21 min of imaging (*n*>159 cells for the ancestor and *n*>148 cells for the proline-evolved population). Horizontal lines represent means.

During growth in DM2500+0.5 M sucrose, the mean growth rate of So0.5-1 cells was 32% higher than that of the ancestor (*p*<10^−4^, two-tailed t-test) while Pr0.5-1 cells grew similarly to the ancestor (~3% difference) (Fig. 4B), consistent with the fitness advantage of So0.5-1 (Fig. 3B) and neutral behavior of Pr0.5-1 (Fig. 3E) relative to the ancestor in high sucrose concentrations. When grown with 0.125 M proline as the sole carbon source, the mean growth rate of Pr0.5-1 cells was 23% higher than that of the ancestor (*p*<10^−4^, two-tailed t-test; Fig. 4C), again consistent with the fitness increase of Pr0.5-1 in this condition. Collectively, these data demonstrate that the adaptations exhibited by both evolved populations are due at least in part to increased ability to grow rapidly.

### Distinct relationships between cell size and growth rate in sorbitol- and proline-evolved populations

Some halophilic species have distinct, atypical morphologies that depend on the osmolarity of their environment (1). Moreover, it has long been recognized that the mean cell volume and growth rate of *E. coli* cells are strongly correlated when growth rate is modulated via the nutrient content of the medium (29, 30), commonly referred to as the Growth Law (31). However, volume varies less strongly with temperature-induced growth rate changes (29), indicating that the specific mechanism of growth rate change is important for determining cell size (32). Given that the ancestor and the sorbitol- and proline-evolved populations exhibited different growth-rate behaviors (Fig. 4), we used these data to query whether cellular dimensions during steady-state exponential growth varied in a manner predictable by nutrient-induced changes in growth rate. The lower growth rate of the ancestor in DM2500+0.5 M sucrose as compared with DM2500 (Fig. 4A, B) was coupled to a decrease in mean cell size (Fig. 5A, B), as expected from the Growth Law. However, while the growth rate of the ancestor in DM+0.125 M proline was higher than DM2500+0.5 M sucrose (Fig. 4B,C), mean cell size was lower in DM+0.125 M proline than in DM2500+0.5 M sucrose (Fig. 5A,B), suggesting more complex relationships among osmolarity, growth rate, cell size than those due to nutrients alone.

**Figure 5:**
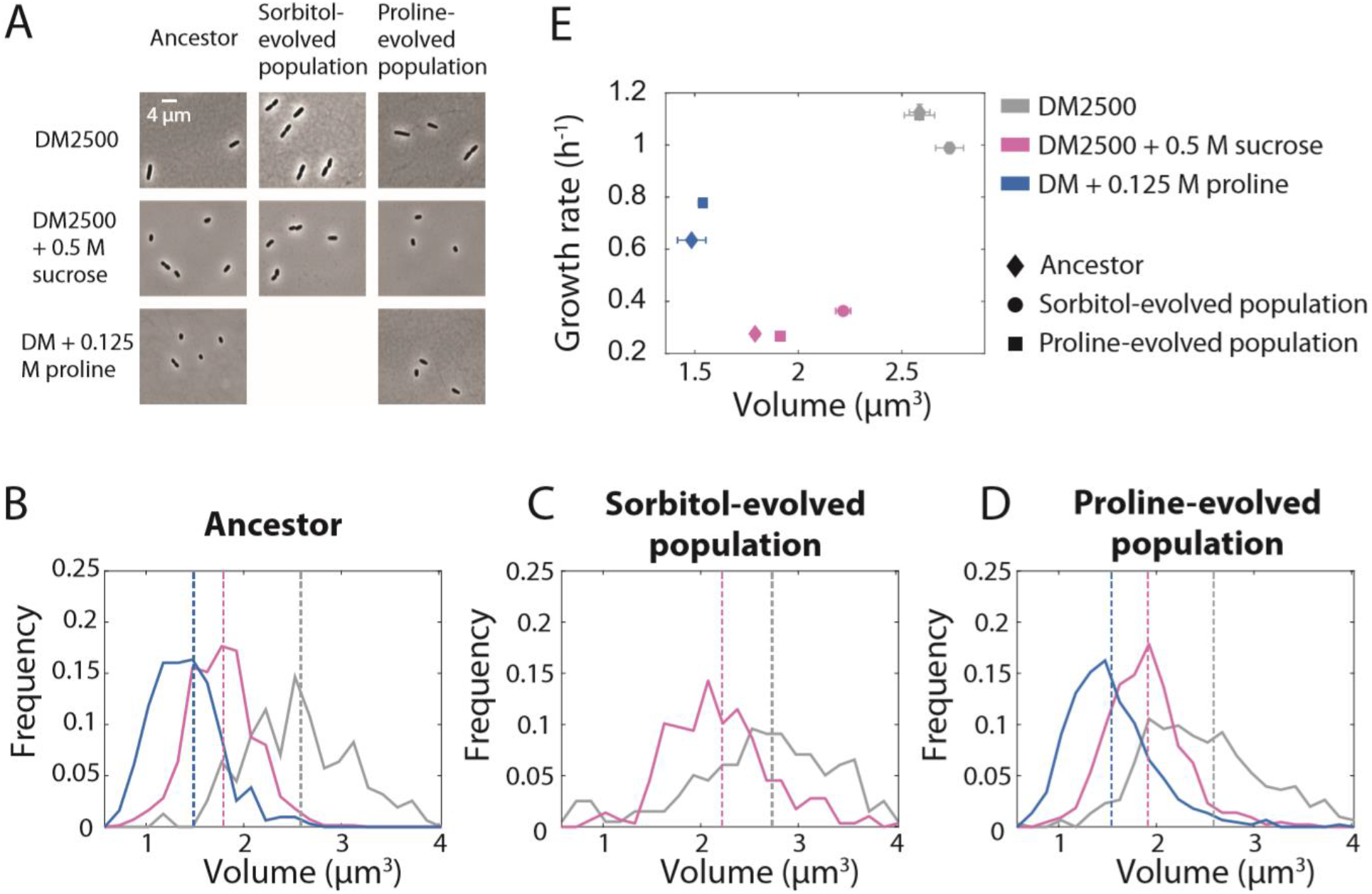
Cell size does not scale with growth rate across media with different osmolytes. A) Typical images of single cells during exponential growth from the ancestor and So0.5-1 and Pr0.5-1 populations in DM2500, DM2500+0.5 M sucrose, and DM+0.125 M proline (*n*>160 cells were imaged for each population). B-D) Normalized histograms of cell volumes of the ancestor (B), So0.5-1 (C), and Pr0.5-1 (D) populations grown in DM2500, DM2500+0.5 M sucrose, and DM+0.125 M proline. All three populations are larger in DM2500 than DM2500+0.5 M sucrose, as expected from the Growth Law based on growth rates (Fig. 4A,B), but the ancestor and Pr0.5-1 population were smaller in DM+0.125 M proline than in DM2500+0.5 M sucrose despite the higher growth rate with proline as a carbon source (Fig. 4B,C). Vertical lines represent medians (*n*>160 cells were imaged for each population). E) Comparison of steady-state growth rates and mean cell volumes across populations and growth conditions demonstrates the lack of agreement with the Growth Law across media or population comparisons, which would predict that the points would fall along a single line. Error bars represent the standard error of the mean. Some error bars are not visible because the error is smaller than the size of the marker (*n*>77 cells were imaged for each population).

We next investigated whether cell size changed in the evolved populations in a systematic manner across media. In DM2500, DM2500+0.5 M sucrose, and DM+0.125 M proline, the So0.5-1 and Pr0.5-1 populations behaved in a qualitatively similar manner as the ancestor (Fig. 5A-D), with the largest cells in DM2500 and the smallest cells in DM+0.125 M proline. So0.5-1 cells were wider (*p*<10^−4^, two-tailed t-test) and longer (*p*<10^−4^, two-tailed t-test) than the ancestor in DM2500+0.5 M sucrose (Fig. S3B), as expected based on their relative growth rates. However, in DM2500, So0.5-1 cells were wider (*p*<10^−4^, two-tailed t-test) and slightly shorter (Fig. S6A) than the ancestor such that the mean volume was slightly higher than the ancestor (Fig. S6A) despite a lower growth rate (Fig. 4A). Thus, there are mutations in the So0.5-1 population that affect cell size independent of the Growth Law.

Relative changes in colony forming units (CFUs) for the ancestor and sorbitol-evolved populations were quantitatively in agreement with OD measurements in both DM2500 and DM25 supplemented with 0.5 M sorbitol (Fig. S7A, B), indicating that variations in cell shape and volume did not measurably impact conclusions based on OD. In the Pr0.5-1 population, cells had approximately the same dimensions as the ancestor (*p*=0.99, two-tailed t-test) in DM2500 (Fig. 5E), consistent with their similar growth rates (Fig. 4A), but were only slightly larger than the ancestor in DM+0.125 M proline (Fig. 5E) despite the significantly higher growth rate of the Pr0.5-1 population (Fig. 4C). Taken together, these results indicate that the So0.5-1 population, which has evolved to generally grow better in high osmolarity conditions, has developed growth rate-independent changes in cell size and aspect ratio, and the Pr0.5-1 population, which has evolved to more efficiently utilize proline as a carbon source, maintained the same size as the ancestor even when their growth rates were different.

### Sorbitol-evolved populations developed mutations involved in central growth processes

To determine the genetic changes that could generate the distinct phenotypes of the sorbitol- and proline-evolved populations, we used metagenomic sequencing and analysis (Methods). In the So0.5-1 population, 83.3% of the reads revealed a 9-bp, in-frame insertion (Table S1) in DNA topoisomerase I (*topA*). In high osmolarity conditions, DNA supercoiling levels increase (33), affecting transcription; TopA relieves supercoiling and is upregulated in response to supercoiling (34). The insertion occurs at base 1329460, which lies in the TOPRIM domain that includes part of the catalytic site (35). The second-most prevalent mutation (17.6%) in the So0.5-1 population was a single base-pair deletion at position 1893043 in the *prc* gene (Table 1), which encodes the carboxyterminal protease for the division-specific transpeptidase Penicillin Binding Protein 3. The deletion shifts the reading frame at position 351 out of 2049, which likely disrupts Prc activity, suggesting that deletion of *prc* could be beneficial in high osmolarity conditions in this background. Previous work showed that Δ*prc* cells are rounder than wild-type cells (36), providing a potential explanation for So0.5-1 cells being wider and shorter than the ancestor in DM2500 (Fig. S3A).

By sequencing the *topA* and *prc* genes in 20 isolates from the So0.5-1 population, we found that 85% possessed the *topA* mutation, consistent with the metagenomics data. All isolates with the *prc* mutation (15%) also possessed the *topA* mutation, indicating that the *prc* allele arose from within the *topA* mutant population. To assess the behavior of these isolates at high osmolarity, we grew a representative with each evolved allele combination in DM2500 and DM25 with 0.5 M sorbitol. When comparing the growth curves of an isolate containing either the *topA* mutation alone or one with both the *topA* and *prc* mutations to the ancestor, we observed that both isolates clearly grew better in DM25+0.5 M sorbitol but worse in DM2500 (Fig. 6A,B). Thus, both mutants have the same qualitative pattern of fitness as the So0.5-1 population.

**Figure 6:**
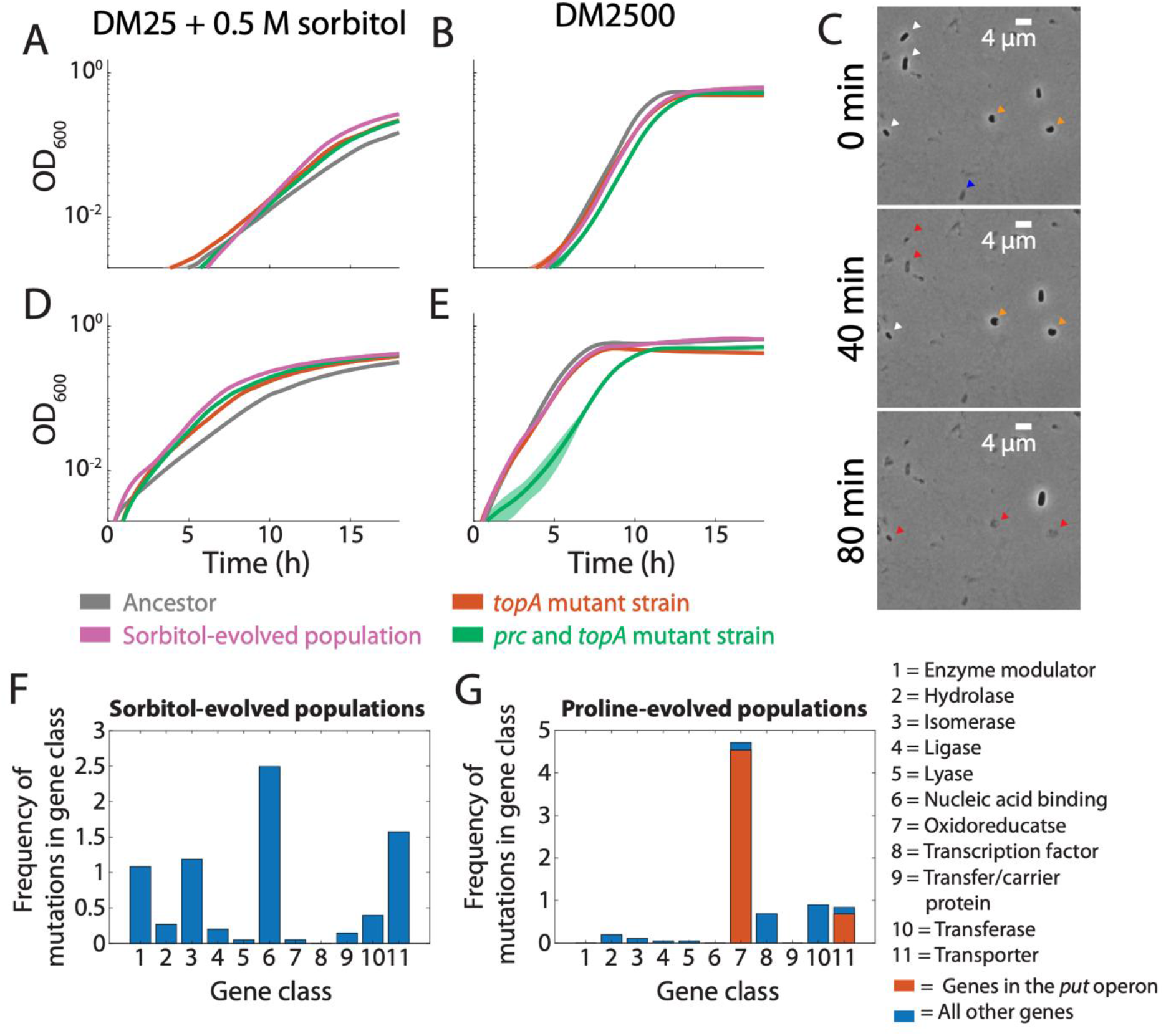
Isolates of the sorbitol-evolved population exhibit fitness tradeoffs consistent with continued adaptation to high osmolarity. A,B) An isolate that contained a mutation in *topA* alone (red) and an isolate that also had a mutation in *prc* (green) had faster growth than the ancestor in DM25+0.5 M sorbitol (A) but slower growth in DM2500 (B). Lines are averages of *n*=6 growth curves, with shaded regions representing standard error of the mean (SEM). The growth curve of the *prc topA* isolate was consistently below that of the *topA* isolate. The SEM is small at some time points and is partially obscured by the lines. C) Time-lapse movie of outgrowth in DM2500 of stationary-phase *prc topA* cells previously grown in DM2500 revealed cells that had already lysed when imaging commenced (blue arrowhead), along with normal cells (white arrowheads) and rounded cells (orange arrowheads) that eventually lysed (red arrowheads). The SEM is small at some time points and is partially obscured by the lines. D,E) The cultures in (A,B) were passaged one additional time in the same media to quantify changes to growth behaviors when the stationary phase prior to growth curve measurement was reached in a growth environment more similar to the passage condition. They were grown in either DM25+0.5 M sorbitol (D) or DM2500 (E). Lines are averages of *n*=6 growth curves, with shaded regions representing SEM. By contrast to (A), the growth curve of the *prc topA* strain was consistently above that of the *topA* strain in DM25+0.5 M sorbitol (D), but substantially lower after an additional passage in DM2500 (E) than in (B). F,G) Genes with mutations identified at >5% prevalence in one of the four So0.5 and Pr0.5 evolved populations were classified by PANTHER Protein Class using PantherDB (http://pantherdb.org)(54). The contribution of each mutation to the histogram was scaled by its prevalence in the population; the orange bars in (G) represent the subset of mutations in the *put* operon. Mutations represented a wide range of functional categories in sorbitol-evolved populations (F), while mutations in the proline-evolved populations were predominantly related to proline utilization. In the four So0.5 populations, 19 of the 44 genes did not have a classification; those genes accounted for 40% when scaled by prevalence. In the four Pr0.5 populations, 25 of the 50 genes did not have a classification; those genes accounted for 34.3% when scaled by prevalence.

However, at lower osmolarity, the isolate containing the *prc* and *topA* mutations grew much worse than either the isolate containing only the *topA* mutation or the evolved So0.5-1 population (Fig. 6B). Moreover, the *topA* and *prc topA* isolates did not grow as well as the So0.5-1 population in the high osmolarity condition (Fig. 6A). Imaging of *prc topA* cells during outgrowth from stationary phase in DM2500 revealed occasional round cells, rod-shaped cells that round up, and lysis (Fig. 6C), suggesting difficulties in surviving and/or emerging from stationary phase. Interestingly, *prc topA* cells occasionally behaved similarly to spheroplasts (37) and grew as large blobs (Movie S1). Thus, slower initial growth of the *prc* <u>*topA*</u> isolate at low osmolarity is likely due to increased cell death during stationary phase or outgrowth.

By contrast, the *topA* strain exhibited better growth in DM25+0.5 M sorbitol than the *prc topA* strain (Fig. 6A), leading us to wonder how the *prc topA* mutation was selected for in the 0.5 M sorbitol passaging medium. Given the imaging data above, we hypothesized that the passaging of strains first in DM25 (low osmolarity) in our fitness assays artificially reduced the fitness of the *prc topA* strain compared to the repeated passages at high osmolarity in our evolution experiment. To test this hypothesis, we passaged the isolated strains and evolved population in DM25+0.5 M sorbitol once before measuring growth curves in DM25+0.5 M sorbitol. With the extra high osmolarity passage, the *prc topA* strain now grew slightly better than the *topA* strain (Fig. 6D), providing evidence for the selective advantage of the *prc* mutation in conditions similar to those during repeated passaging. By contrast, a passage in DM2500 before measuring growth curves in DM2500 further exacerbated the fitness defects of *prc topA* in low osmolarity (Fig. 6E).

Metagenomic sequencing of the three other populations passaged in 0.5 M sorbitol also revealed high prevalence mutations with diverse sets of potential functional consequences (Table 1, Fig. 6F). An 84-bp deletion at the 3’ end of *topA* in population 4 replaced the last 7 amino acids with 11 different amino acids, further implicating TopA in fitness at high sorbitol concentrations (Fig. 2C). Populations So0.5-2 and So0.5-4 had mutations in the transcription elongation factor *nusA* present at >50%, and So0.5-1 had a low prevalence mutation in the transcription termination factor *rho*, further demonstrating the importance of transcriptional regulation and NusA in particular in high osmolarity conditions. These observations suggest that the supercoiling induced in high osmolarity conditions is a key element of the stress induced in high osmolarity conditions and that many different proteins whose function is closely tied to DNA structure can be readily mutated to increase fitness in high osmolarity conditions. Across all four populations, there are >10 genes (Table 1) involved in transport that acquired mutations, demonstrating that transport-related genes are a large and key target for selection in high osmolarity. Two of these genes are previously of unknown function; their appearance in sorbitol-evolved populations suggests roles in growth at high osmolarity. *yfdV*, which is predicted to encode a transporter (38), had a 9-bp IS*1* insertion at 21% prevalence in So0.5-3. *ycfT*, which is predicted to encode an inner membrane protein (39) but is not predicted to be a transporter (38), had a 9-bp IS*1* insertion in So0.5-4 at 65% prevalence. Based on the observation that mutations in these genes appeared in evolved populations with an increased prevalence of mutations in transport-related genes, we speculate that that *ycfT* interacts with transporters and that *yfdV* is in fact a transporter. Mutations in central metabolism (*sucA*), cell-envelope synthesis (*prc*, *ftsW*, and *slyB*), mechanosensation (*mscL*), and phosphate regulation (*phoR*, *pstS*) also appeared with high prevalence. Taken together, these observations show that high sorbitol concentration can select for mutations affecting a wide range of cellular processes, with enzymes that interact with DNA and transporters being the two most frequently targeted classes of proteins (Fig. 6F).

### Evolution in proline predominantly selects for mutations in genes regulating proline metabolism

To determine whether the Pr0.5-1 population developed similar mutations as the sorbitol-evolved population in response to osmolyte pressure, we again used metagenomic sequencing. We identified a fixed (100%) C→T mutation in *putA* that resulted in a glycine to arginine substitution at the 7^th^ position of the reading frame (Table S1). This mutation is in a DNA-binding domain that functions as a repressor of *putA*/*putP* (40), the genes responsible for growth on proline (41). The PutA enzyme oxidizes proline to glutamate so that *E. coli* can utilize it as a carbon source, and *putP* encodes a membrane-bound proline: Na+ symporter (42, 43). Based on our growth data with proline (Fig. 2B, D), this mutation very likely inhibits binding of the repressor domain, which results in upregulation of the genes responsible for proline catabolism.

Metagenomic sequencing of the other three populations evolved in 0.5 M proline also showed much less diversity in mutated genes as compared with the sorbitol-evolved populations (Table S1, Fig. 6G). Populations Pr0.5-1, 2, and 4 had a mutation in *putA* present in 92-100% of the population; for population 3, there were multiple mutations at 69% and 24% prevalence (Table 1). There was also an *arcB* mutation in population 2 at 15% and an *arcA* mutation in population 4 at 35%; *arcAB* encode a two-component system that regulates proline metabolism. Thus, unlike evolution in sorbitol, the readily targetable proline metabolism genes provide large growth benefits on proline and hence mutations specific to this advantage are more often selected for.

## Discussion

Here, we aimed to investigate how osmolarity affects cell growth and morphology by examining the genetic and physiological changes that occurred during evolution in a variety of environments. Comparisons of these changes across combinations and concentrations of osmolytes provided insight into the nature of stress presented by a given evolution condition and the mechanisms cells can utilize to optimize growth in a new environment. Although the evolved populations in 0.5 M sorbitol and 0.5 M proline clearly had higher fitness than the ancestor in their respective growth environments, our study revealed that the selective pressures imposed by these two osmolytes were distinct. Evolution in sorbitol resulted in general adaptation to high osmolarity, at the expense of growth at lower osmolarity (Fig. 3B-D). By contrast, evolution in proline resulted in improved proline metabolism, without impacting the response to any level of increased osmolarity (Fig. 3E). Further studies are necessary to ascertain whether it is generally the case that exposure to molecules that directly function in cellular metabolism and osmoregulation, as is the case with proline, will primarily result in genetic perturbations to a specific pathway first. These findings illustrate the complex nature of osmotic stress, particularly for osmolytes that can also be used as carbon sources.

Our data strongly indicate that *topA* plays a major role during growth in high osmolarity conditions. The *topA* mutations that occurred are likely adaptive, given that *topA* exhibited the highest prevalence of mutation during passaging in sorbitol and the increased growth rate of the So0.5-1 population in many different high osmolarity conditions relative to the ancestor (Fig. 3C, D, S4). The So0.5-4 population, which has higher fitness than the ancestor in 0.5 M sorbitol (Fig. 2A,C), also contained a *topA* mutation at 25% prevalence. During growth at high osmolarity, increased DNA supercoiling is thought to be beneficial because genes that respond to osmotic stress such as *proP*, which encodes a glycine betaine/proline transport system, are positively regulated by negative supercoiling (33, 44, 45). Thus, relieving of the active negative supercoiling by TopA would waste energy and result in less effective activation of genes necessary for survival at high osmolarity. Single amino-acid substitutions in *topA* that increase negative supercoiling have been previously identified in long-term evolution experiments in DM25 (46, 47). The three amino-acid insertion in the active site of *topA* in the So0.5-1 population likely reduces its activity to an even greater degree, which would confer a fitness advantage at high osmolarities but could negatively impact cells at low osmolarities due to the deleterious effects of excessive negative supercoiling, consistent with our population fitness measurements (Fig. 3D). The lower fitness of the So0.5-1 population relative to the ancestor in media with low (0.125 M or less) sorbitol or sucrose concentrations indicates that the optimal osmolarity for growth of this population has increased. These findings raise the interesting possibility that tuning TopA activity could continuously shift the optimal osmolarity, which may have important implications in biotechnology.

In a seminal long-term experiment in which *E. coli* was passaged daily into fresh minimal medium for more than 50,000 generations, the fitness of the replicate populations (defined as the ability to outcompete the ancestor in co-culture) first increased rapidly for ~2,000 generations (33), then continued to increase at a much slower rate (34). Over the first 2,000 generations the lag time decreased while the growth rate increased; overall the fitness benefit from the growth rate change was higher (35), consistent with the increased growth rates of our evolved populations in their passaging medium (Fig. 4). After ~10,000 generations, average cell volume had also increased substantially (17, 33, 35, 36). The evolved populations from that experiment exhibited a stepwise, allometric correlation between cell size and growth rate (21), in agreement with previous studies of the effects of nutrients on cell size (31, 32). In a separate experiment, clones isolated from 115 *E. coli* populations evolved for ~2,000 generations at 42 °C had substituted a total of more than 1,300 mutations (37); while almost no mutations were shared between replicates, there was strong convergence at the gene, operon, and functional complex level, including a large fraction of mutations in genes associated with cell-wall synthesis (37). Thus, it appears that specific pathways may evolve in response to a global environmental stress, although it is unknown whether the resulting mutations will be more or less widespread during evolution under other selective pressures. Our results generally support qualitative phenotypic differences among populations evolved in sorbitol and proline (Fig. 2-5), indicating that bacterial cells experience osmolyte-specific fitness costs during osmotic stress. The action of mutations that confer increased fitness in one osmolyte can extend in a complex manner to other osmolytes, as evidenced by the similar advantages in the sorbitol-evolved population relative to the ancestor in both sorbitol and sucrose (Fig. 3B-D). Interestingly, none of the mutations that we observed in sorbitol-evolved populations occurred in genes previously identified as being highly up- or down-regulated in *E. coli* during growth in media supplemented with 0.7 M sorbitol (24). This observation demonstrates the power of laboratory evolution experiments: the fact that expression of a particular gene is integral to a stress response does not necessarily signify that it is a readily mutable target for selection, and some of the targets for adaptive mutations may not normally be associated with the stress. While the genes that acquired mutations in the four sorbitol-evolved populations encode proteins involved in a wide range of functions, transport and transcription are clearly key targets for beneficial mutations at high osmolarity (Fig. 6F, Table 1).

It remains unclear why no adaptation was observed to any of the other osmolyte passage conditions, when generally improved fitness at high osmolarity was readily evolvable in sorbitol. One possibility is that the passage concentrations of sucrose (0.1 and 0.2 M) did not exert enough stress to select for beneficial mutations; consistent with this conclusion, there was no discernable change in growth during passaging in 0.25 M sorbitol. However, this argument does not explain the lack of adaptation in all 0.5 M and 0.75 M glycine betaine populations. Notably, the only conditions in which we observed adaptation are also those in which the osmolyte can be used as a carbon source, thus leading to a higher carrying capacity. While each passage involves a similar number of generations due to the constant degree of back-dilution, the increased population sizes during growth in proline and sorbitol allow for exploration of a larger mutational space per generation. Thus, it is possible that adaptation in the other osmolytes could be accelerated by increasing the amount of glucose to bring the yield in line with that of the sorbitol and proline populations; it is also possible that more extensive passaging or increasing the osmolyte concentration would spur adaptation. Regardless, metabolism of sorbitol did not preclude the selection of mutations that generally increase fitness at high osmolarity, as evidenced by growth in sucrose (Fig. 3B, D).

In combination with new tools for phenotypic quantification and inexpensive sequencing, long-term evolution experiments have tremendous potential for providing unique molecular and physiological insight into how bacteria adapt to new environments. Here, we uncovered general and specific factors regulating growth at higher osmolarities that had previously not been associated with such conditions. Our data demonstrate that *E. coli* has access to a wide variety of mutations that improve fitness by mitigating the broad detrimental effects of high osmolarity. Determining the extent to which the same is true for other stressful environments should shed light on the general nature of cellular adaptation, motivating future evolution experiments with other perturbations and in other organisms.

## Methods and Materials

### Passaging and evolution of cultures

*E. coli* TC1407 was grown overnight in DM25 (48, 49) at 37 °C and then used to inoculate a deep-well 96-well plate (catalog number 780285, Greiner Bio-One, Germany) with media of various osmolytes and concentrations. We chose 10 experimental conditions with higher osmolarity, one DM25 condition, and one blank control. Our experimental conditions were selected to cover a wide range of osmolarities and osmolytes: 0.5 M and 1.0 M glycine betaine, 0.5 M and 0.75 M proline, 0.25 M and 0.5 M sorbitol, 0.1 M and 0.2 M sucrose, and M and 0.2 M NaCl. We passaged four replicate wells of each condition, in addition to four blank wells to control for contamination (Fig. 1A). Each column of the 96-well plate corresponded to a separate condition. To initialize the experiment, 10 μL of the overnight cultures and 1 mL osmolyte medium were added to each well in a 96-well plate. The plate was covered with a Breathe-Easy membrane (catalog number 9123-6100, USA Scientific, Florida) and incubated at 37 °C overnight with shaking. Each day thereafter, for a total of 38 days, 10 μL of the overnight cultures were transferred to wells of a new 96-well plate with 1 mL of the appropriate media. The 100-fold dilutions yielded 6.6 generations of growth per day as long as the population reached stationary phase.

### Measurement of population growth metrics

We measured population-level growth metrics using an Epoch 2 Microplate Spectrophotometer (Biotek Instruments, Vermont). A 200-μL 96-well plate filled with DM25 medium was inoculated from glycerol stocks and grown for 24 h, followed by two cycles of 1:200 dilutions into fresh DM25 and 24 h of growth. The resulting cultures were used to inoculate 200-μL cultures in a 96-well plate with a 1:200 dilution into DM25 or DM2500 medium with osmolytes added as indicated. The plate reader went through 15-min cycles of incubation at 37 °C, shaking linearly for 145 s, and then absorbance measurements (wavelength 600 nm, 25 flashes, 2-ms settle between flashes). Growth curves were performed for 24-72 h, which allowed most populations to reach stationary phase. CFUs/mL were measured by serially diluting 10 μL of culture from a plate reader experiment, plating 100 μL of each dilution, and counting colonies after 16 h of growth.

### Single-cell imaging

Cells were imaged on a Nikon Eclipse Ti-E inverted fluorescence microscope with a 100× (NA 1.40) oil-immersion objective (Nikon Instruments). Images were collected on a DU885 electron-multiplying CCD camera (Andor Technology) or a Neo sCMOS camera (Andor Technology) using μManager v. 1.4 (50). Cells were maintained at 37 °C during imaging with an active-control environmental chamber (Haison Technology). For experiments conducted on agarose pads, 1 μL of cells was spotted onto a pad of 1% agarose + medium as noted.

### Image analysis

The MATLAB (MathWorks, Natick, MA, USA) image processing code *Morphometrics* (51) was used to segment cells and to identify cell outlines from phase-contrast microscopy images. A local coordinate system was generated for each cell outline using a method adapted from *MicrobeTracker* (52). Cell widths were calculated by averaging the distances between contour points perpendicular to the cell midline, excluding contour points within the poles and sites of septation. Cell length was calculated as the length of the midline from pole to pole. Cell volume was estimated from width and length measurements by approximating cells as a pill shape with volume 2*πR*^2^(*L*−2*R*) + 4/3*πR*^3^, where *R* is half the cell width and *L* is the cell length. Cellular dimensions were quantified by averaging single-cell results across a population. Single-cell growth rates were calculated using the formula 1/Δ*t* ln(*V*(*t*+*Δt*)/*V*(*t*)), where Δ*t* is the time between images and *V*(*t*+*Δt*) and *V*(*t*) are the cell volumes at times *t*+Δ*t* and *t*, respectively. See figure legends for the number of cells analyzed (*n*) and error bar definitions.

### Whole-genome sequencing

Cultures were grown from a glycerol −80 °C frozen stock in LB overnight. Genomic DNA was extracted from these overnight cultures using the DNeasy UltraClean Microbial Kit (Qiagen), and prepared for sequencing using the Nextera library kit. Sequencing was performed on an Illumina NovaSeq 6000 at ~100X coverage. Read mapping and variant calling was accomplished using BreSeq v. 0.33.0 (53).

### Data availability

All data used in this manuscript are growth curves, time-lapse microscopy images, or whole-genome sequencing. All data are available upon request from the corresponding author.

## Acknowledgments

We thank Russell Monds for helpful discussions on the design of the evolution experiments, and Lisa Willis and Heidi Arjes for careful readings of the manuscript. Funding was provided by NSF CAREER Award MCB-1149328 and the Allen Discovery Center at Stanford on Systems Modeling of Infection (to K.C.H.), and a Stanford Graduate Fellowship and an NSF Graduate Research Fellowship (to S.C.). M.A.D. and E.L. were supported by the Stanford Bioengineering Research Experience for Undergraduates program. This work was also supported in part by the National Science Foundation under grant PHYS-1066293 and the hospitality of the Aspen Center for Physics. K.C.H. is a Chan Zuckerberg Biohub Investigator.

