## Supplemental Tables and Figures for "Bacterial evolution in high osmolarity environments"

### 1 Supplementary Tables

| Population | Position | Mutation | Frequency | Annotation | Gene | Description |
|---|---|---|---|---|---|---|
| So0.5-1 | 1329460 | +CAAAA CGAT | 0.833 | coding (41/2598 nt) | *topA* → | DNA topoisomerase I |
| So0.5-1 | 1893043 | Δ1 bp | 0.176 | coding (351/2049 nt) | *prc* ← | carboxy-terminal protease for penicillin-binding protein 3 |
| So0.5-1 | 3871975 | IS1 (–) +8 bp | 0.094 | intergenic (84/+223) | *pstS* ← / ← *glmS* | phosphate transporter subunit/D-fructose-6-phosphate amidotransferase |
| So0.5-1 | 3928330 | +CTGAAG | 0.091 | coding (114/1260 nt) | *rho* → | transcription termination factor Rho |
| So0.5-2 | 101978 | T→A | 0.184 | F258I (TTT→ATT) | *ftsW* → | integral membrane protein involved in stabilizing FtsZ-ring during cell division |
| So0.5-2 | 386758 | IS1 (–) +9 bp | 0.614 | coding (447455/ 1296 nt) | *phoR* → | sensory histidine kinase in two-component regulatory system with PhoB |
| So0.5-2 | 742545 | Δ132 bp | 0.143 | coding (233364/ 1218 nt) | *sucB* → | dihydrolipoamide acetyltransferase |
| So0.5-2 | 3015771 | IS150 (–) +3 bp | 0.162 | coding (11921194/ 1677 nt) | *ECB_02816* → | KpsD protein |
| So0.5-2 | 3251730 | A→G | 0.533 | L344P (CTG→CCG) | *nusA* ← | transcription elongation factor NusA |
| So0.5-2 | 3365992 | G→A | 0.287 | G22S (GGT→AGT) | *mscL* → | large conductance mechanosensitive channel |
| So0.5-2 | 3625458 | T→A | 0.131 | pseudogene (5/ 738 nt) | *yhjQ* ← | b3534; cell-division protein (chromosome partitioning ATPase) |
| So0.5-3 | 16972 | IS150 (–) +3 bp | 0.078 | intergenic (14/514) | *mokC* ← / → *nhaA* | regulatory protein for HokC, overlaps CDS of *hokC*/pH-dependent sodium/proton antiporter |
| So0.5-3 | 1166458 | T→G | 0.079 | intergenic (+38/50) | *acpP* → / → *fabF* | acyl carrier protein/3-oxoacyl-(acyl-carrier-protein) synthase |
| So0.5-3 | 1166459 | T→G | 0.069 | intergenic (+39/49) | *acpP* → / → *fabF* | acyl carrier protein/3-oxoacyl-(acyl-carrier-protein) synthase |

| | | | | | | |
|---|---|---|---|---|---|---|
| So0.5-3 | 1434768 | IS150 (+) +3 bp | 0.107 | coding (112114/ 2640 nt) | *ydbH* → | hypothetical protein |
| So0.5-3 | 1462251 | IS150 (–) +3 bp | 0.275 | intergenic (13/326) | *mokB* ← / → *trg* | regulatory peptide/methyl-accepting chemotaxis protein III, ribose and galactose sensor receptor |
| So0.5-3 | 1697515 | A→T | 0.094 | Q149L (CAG→CTG) | *slyB* → | outer membrane lipoprotein |
| So0.5-3 | 2416047 | C→A | 0.063 | M119I (ATG→ATT) | *emrY* ← | predicted multidrug efflux system |
| So0.5-3 | 2424024 | IS1 (+) +9 bp | 0.21 | coding (382390/ 945 nt) | *yfdV* ← | predicted transporter |
| So0.5-3 | 2813703 | IS150 (+) +3 bp | 0.202 | coding (505507/ 546 nt) | *syd* ← | SecY interacting protein Syd |
| So0.5-3 | 3968420 | G→T | 0.063 | T65K (ACG→AAG) | *rarD* ← | predicted chloramphenicol resistance permease |
| So0.5-3 | 4096974 | A→C | 1 | F200C (TTT→TGT) | *glpF* ← | glycerol facilitator |
| So0.5-3 | 4504909 | T→C | 0.175 | intergenic (+653/ 72) | *insB*25 → / → *ECB_04162* | b4563(b4576); IS1 protein *insB*/hypothetical protein |
| So0.5-4 | 741343 | A→C | 0.579 | E616A (GAA→GCA) | *sucA* → | alpha-ketoglutarate decarboxylase |
| So0.5-4 | 741844 | +TCA | 0.063 | coding (2348/ 2802 nt) | *sucA* → | alpha-ketoglutarate decarboxylase |
| So0.5-4 | 1189282 | IS1 (+) +9 bp | 0.649 | coding (471479/ 1074 nt) | *ycfT* ← | predicted inner membrane protein |
| So0.5-4 | 1331998 | Δ84 bp | 0.25 | coding (2578/2598 nt) | *topA* → | DNA topoisomerase I |
| So0.5-4 | 2647312 | G→A | 0.06 | intergenic (73/ +248) | *kgtP* ← / ← *rrfG* | alpha-ketoglutarate transporter/5S ribosomal RNA |
| So0.5-4 | 3251631 | Δ27 bp | 0.676 | coding (11041130/ 1488 nt) | *nusA* ← | transcription elongation factor NusA |
| So0.5-4 | 3871028 | Δ1 bp | 0.242 | coding (864/ 1041 nt) | *pstS* ← | phosphate transporter subunit |
| Pr0.5-1 | 1095148 | C→T | 1 | G7R (GGG→AGG) | *putA* ← | fused DNA-binding transcriptional regulator/proline dehydrogenase/ pyrroline-5-carboxylate dehydrogenase |
| Pr0.5-1 | 3312628 | T→C | 0.101 | L10P (CTA→CCA) | *argR* → | arginine repressor |

| | | | | | | |
|---|---|---|---|---|---|---|
| Pr0.5-2 | 858245 | C→A | 0.06 | G224G (GGG→GGT) | *moeA* ← | molybdopterin biosynthesis protein |
| Pr0.5-2 | 877515 | C→A | 0.07 | D90Y (GAC→TAC) | *ybjH* ← | hypothetical protein |
| Pr0.5-2 | 1095079 | G→T | 0.92 | H30N (CAC→AAC) | *putA* ← | fused DNA-binding transcriptional regulator/proline dehydrogenase/ pyrroline-5-carboxylate dehydrogenase |
| Pr0.5-2 | 1592657 | C→A | 0.069 | S40* (TCA→TAA) | *ydeJ* → | competence damage-inducible protein A |
| Pr0.5-2 | 1752551 | C→A | 0.078 | I200I (ATC→ATA) | *aroD* → | 3-dehydroquinate dehydratase |
| Pr0.5-2 | 2451950 | C→A | 0.066 | P88T (CCG→ACG) | *yfeD* → | predicted DNA-binding transcriptional regulator |
| Pr0.5-2 | 3287763 | G→T | 0.147 | D166E (GAC→GAA) | *arcB* ← | hybrid sensory histidine kinase in two-component regulatory system with ArcA |
| Pr0.5-2 | 3625458 | T→A | 0.239 | pseudogene (5/ 738 nt) | *yhjQ* ← | b3534; cell-division protein (chromosome partitioning ATPase) |
| Pr0.5-2 | 4586040 | G→T | 0.069 | E273* (GAA→TAA) | *yjjN* → | predicted oxidoreductase, Zn-dependent and NAD(P)binding |
| Pr0.5-3 | 1045926 | T→G | 0.28 | I350L (ATC→CTC) | *yccW* ← | predicted methyltransferase |
| Pr0.5-3 | 1094443 | G→C | 0.692 | P242A (CCG→GCG) | *putA* ← | fused DNA-binding transcriptional regulator/proline dehydrogenase/ pyrroline-5-carboxylate dehydrogenase |
| Pr0.5-3 | 1094878 | C→T | 0.242 | A97T (GCG→ACG) | *putA* ← | fused DNA-binding transcriptional regulator/proline dehydrogenase/ pyrroline-5-carboxylate dehydrogenase |
| Pr0.5-3 | 1095204 | G→A | 0.685 | intergenic (38/385) | *putA* ← / → *putP* | fused DNA-binding transcriptional regulator/proline dehydrogenase/ pyrroline-5-carboxylate dehydrogenase/ proline:sodium symporter |
| Pr0.5-3 | 2258855 | IS186 (–) +6 bp | 0.314 | coding (586591/ 651 nt) | *alkB* ← | oxidative demethylase of N1-methyladenine or N3-methylcytosine DNA lesions |

| | | | | | | |
|---|---|---|---|---|---|---|
| Pr0.5-3 | 3312631 | Δ9 bp | 0.095 | coding (3240/ 471 nt) | *argR* → | arginine repressor |
| Pr0.5-3 | 3356398 | T→A | 0.162 | noncoding (270/ 1542 nt) | *rrsD* ← | 16S ribosomal RNA |
| Pr0.5-3 | 3463319 | T→G | 0.085 | I412L (ATC→CTC) | *envZ* ← | osmolarity sensor protein |
| Pr0.5-3 | 3602185 | T→G | 0.069 | L118V (TTA→GTA) ‡ | *yhjC* → | predicted DNA-binding transcriptional regulator |
| Pr0.5-3 | 3602186 | T→G | 0.06 | L118* (TTA→TGA) ‡ | *yhjC* → | predicted DNA-binding transcriptional regulator |
| Pr0.5-3 | 3625458 | T→A | 0.121 | pseudogene (5/ 738 nt) | *yhjQ* ← | b3534; cell division protein (chromosome partitioning ATPase) |
| Pr0.5-3 | 4141540 | IS1 (+) +8 bp | 0.084 | coding (613620/ 705 nt) | *yijC* → | DNA-binding transcriptional repressor |
| Pr0.5-4 | 1094872 | A→C | 1 | Y99D (TAT→GAT) | *putA* ← | fused DNA-binding transcriptional regulator/proline dehydrogenase/ pyrroline5carboxylate dehydrogenase |
| Pr0.5-4 | 1322850 | C→A | 1 | A25A (GCC→GCA) | *insA*11 → | IS1 protein InsA |
| Pr0.5-4 | 3127587 | G→T | 0.082 | intergenic (+226/ +567) | *yqiK* → / ← *rfaE* | hypothetical protein/fused heptose 7-phosphate kinase/heptose 1-phosphate adenyltransferase |
| Pr0.5-4 | 4628175 | C→T | 0.348 | D98N (GAT→AAT) | *arcA* ← | DNA-binding response regulator in two-component regulatory system with ArcB or CpxA |
| Pr0.5-4 | 4358156 | +A | 0.137 | coding (328/396 nt) | *frdC* ← | fumarate reductase subunit C |

**Table S1: Mutations identified via metagenomic sequencing of the So0.5 and Pr0.5 populations.** Mutations that were present at <5% were not included in the table.

**Supplementary Figure Legend**

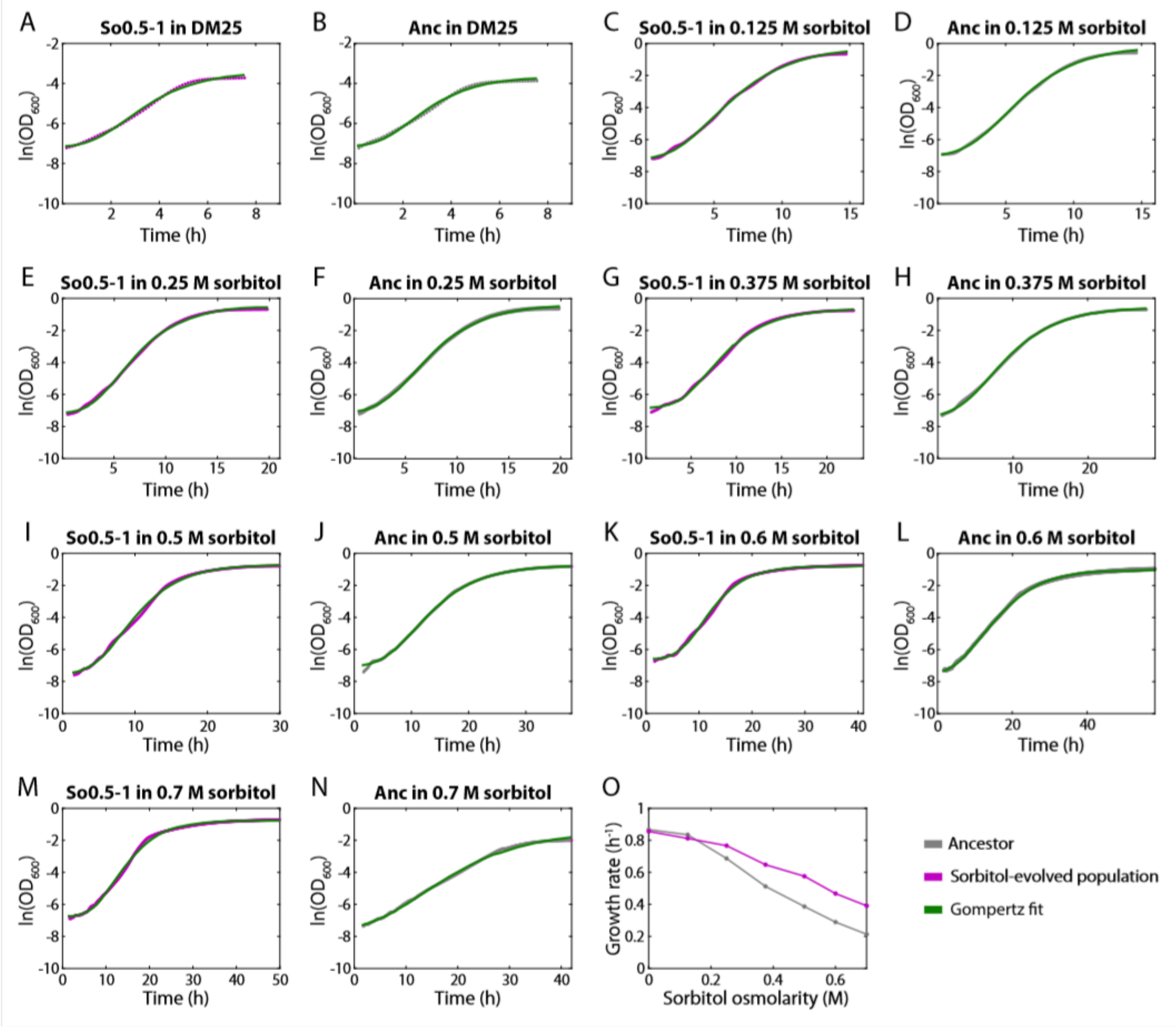


**Supplementary Figure 1: Growth curves of evolved populations and the ancestor in the evolution passage condition.**

Growth curves of the populations evolved in DM25 alone (A), DM25 supplemented with 0.5 M (B) or 0.75 M (C) glycine betaine, 0.25 M sorbitol (D), 0.1 M NaCl (E), 0.1 M (F) or 0.2 M (G) sucrose, or 0.75 M proline (H) was similar to the ancestor in the same medium. Note that some cultures were difficult to revive from frozen stocks after passaging, including one of the DM25 populations (A), 3 of the 4 populations in 0.5 M glycine betaine (B), one of the

0.25 M sorbitol populations (D), all NaCl populations except one in 0.1 M (E), one of the 0.2 M sucrose populations (G), and 3 of the 4 populations in 0.75 M proline (H). In the case of 0.75 M proline, this loss may be due to the lack of saturation after 24 h (Fig. 1F); for NaCl, loss may reflect a reduction in viability during freezing. Growth curves are averages of $n$=3 replicates for each evolved population and averages of $n$=6 replicates for the ancestor, with shaded regions representing the standard error of the mean (SEM). The SEM is small at some time points and is partially obscured by the line.

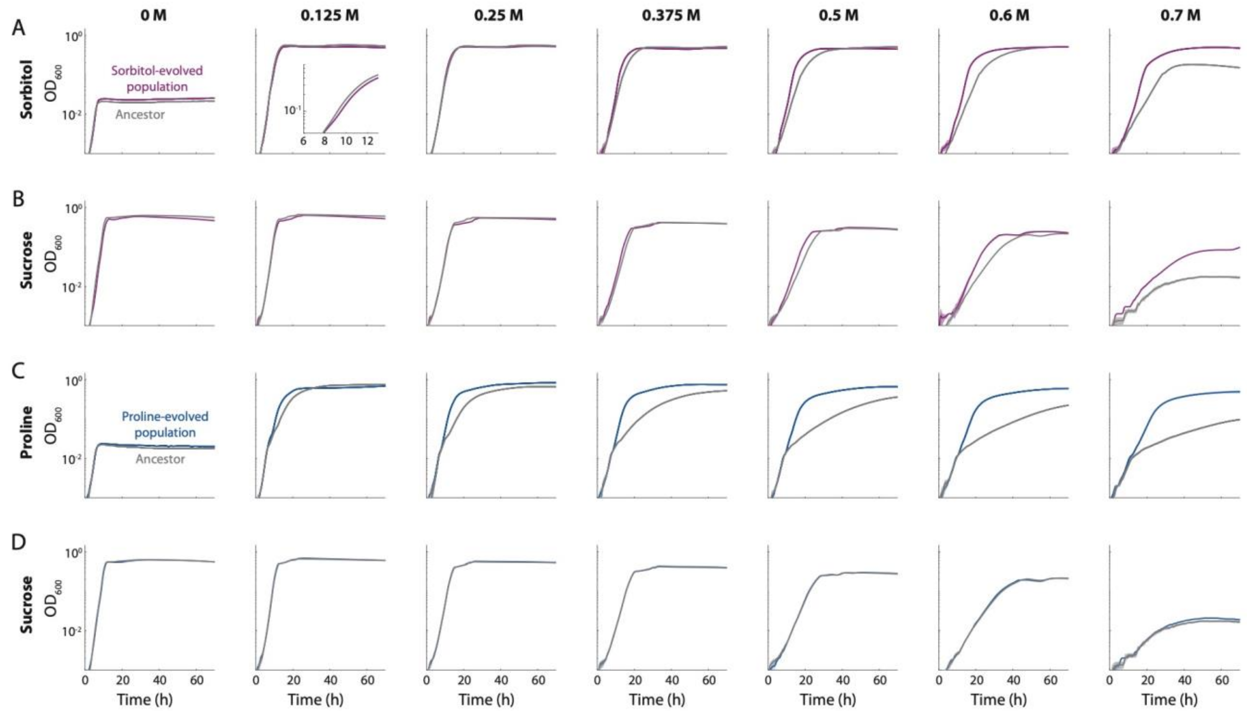


**Supplementary Figure 2: Growth curves for the evolved populations and the ancestor across concentrations of the osmolyte used for passaging and sucrose.**

A) Growth curves of the ancestor (gray) and So0.5-1 population (purple) grown in DM25 with 0 M, 0.125 M, 0.25 M, 0.375 M, 0.5 M, 0.6 M, or 0.7 M sorbitol added. Inset for 0.125 M is a zoom-in demonstrating a slight fitness disadvantage for the sorbitol-evolved population. Growth curves are averages of *n*=4 replicates, with shaded regions representing the standard error of the mean (SEM). The SEM is small at some time points and is partially obscured by the lines.

B) Growth curves of the ancestor (gray) and So0.5-1 population (purple) grown in DM2500 with 0 M, 0.125 M, 0.25 M, 0.375 M, 0.5 M, 0.6 M, or 0.7 M sucrose added. Growth curves are averages of *n*=4 replicates, with shaded regions representing the SEM. The SEM is small at some time points and is partially obscured by the lines.

C) Growth curves of the ancestor (gray) and Pr0.5-1 population (blue) grown in DM25 with 0 M, 0.125 M, 0.25 M, 0.375 M, 0.5 M, 0.6 M, or 0.7 M proline added. Growth curves are averages of $n=4$ replicates, with shaded regions representing the SEM. The SEM is small at some time points and is partially obscured by the lines.

D) Growth curves of the ancestor (gray) and Pr0.5-1 population (blue) grown in DM2500 with 0 M, 0.125 M, 0.25 M, 0.375 M, 0.5 M, 0.6 M, or 0.7 M sucrose added. Growth curves are averages of $n=4$ replicates, with shaded regions representing the SEM. The SEM is small at some time points and is partially obscured by the lines.

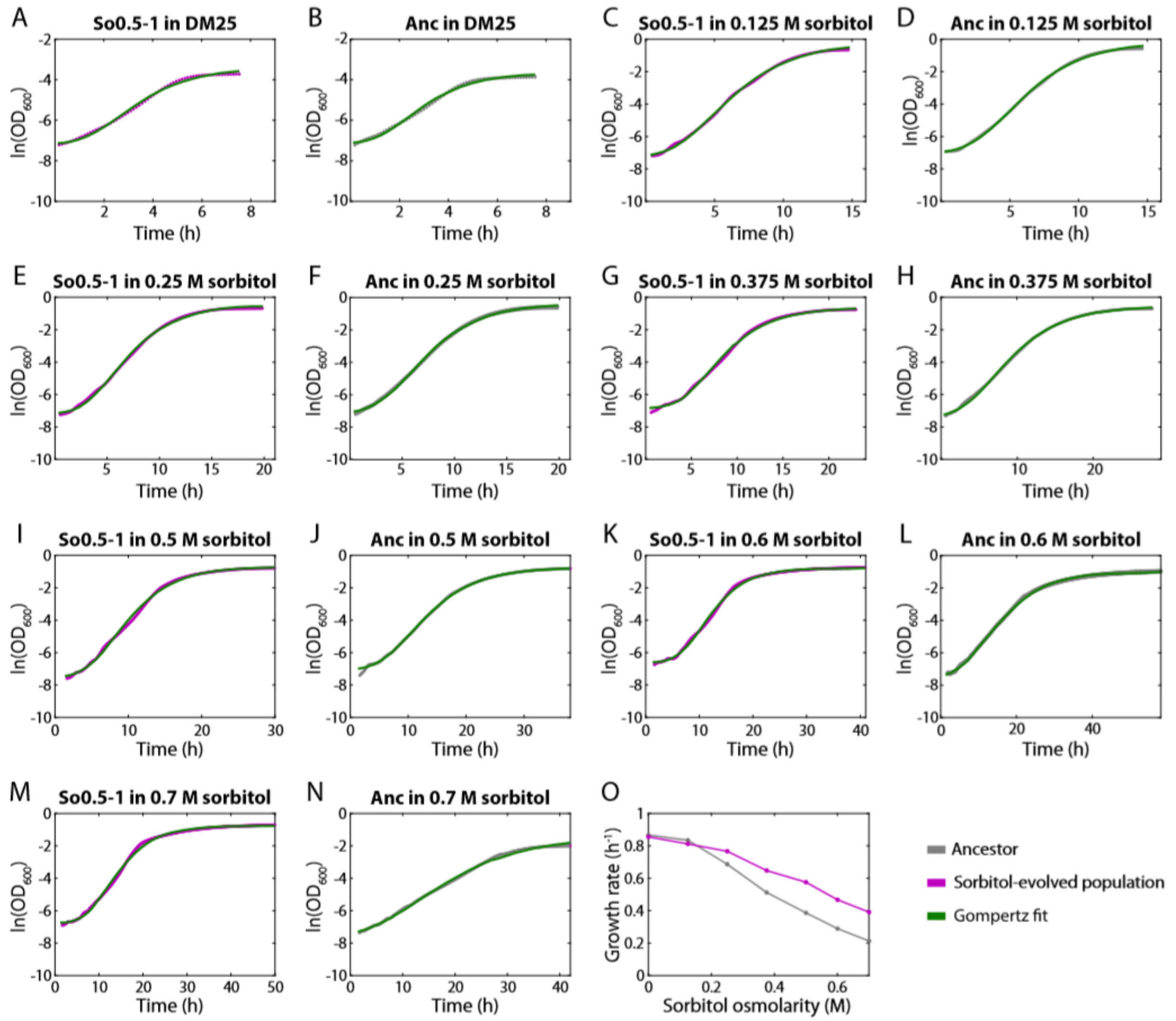


**Supplementary Figure 3: A Gompertz relation provides a good fit to growth curves of the sorbitol-evolved population and the ancestor across all sorbitol concentrations.** Growth curves (gray or magenta) and overlaid Gompertz fits (green) of the So0.5-1 population and ancestor (Anc) grown in DM25 supplemented with 0 M (A, B), 0.125 M (C, D), 0.25 M (E, F), 0.375 M (G, H), 0.5 M (I, J), 0.6 M (K, L), or 0.7 M (M, N) sorbitol. (O) The maximum growth rate extracted from the Gompertz fits of the ancestor and the sorbitol-evolved populations across concentrations of sorbitol was consistent with growth-rate measurements in Fig. 3D.

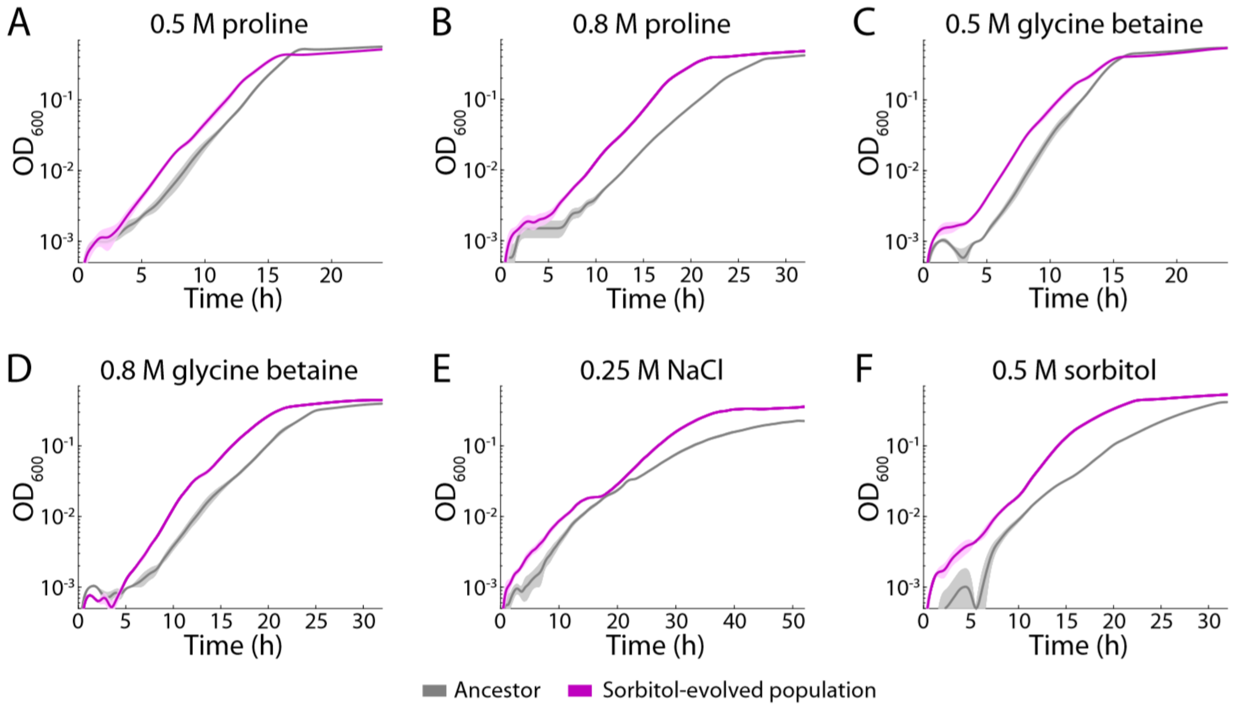


**Supplementary Figure 4: The sorbitol-evolved population is generally more fit than the ancestor at high osmolarity regardless of the osmolyte.**

Growth curves of the ancestor (gray) and So0.5-1 population (purple) grown in DM2500 supplemented with 0.5 M (A) or 0.8 M (B) proline, 0.5 M (C) or 0.8M (D) glycine betaine, 0.25 M NaCl (E), and 0.5 M sorbitol (F). Growth curves are averages of *n*=3 replicates, with shaded regions representing the standard error of the mean (SEM). The SEM is small at some time points and is partially obscured by the lines.

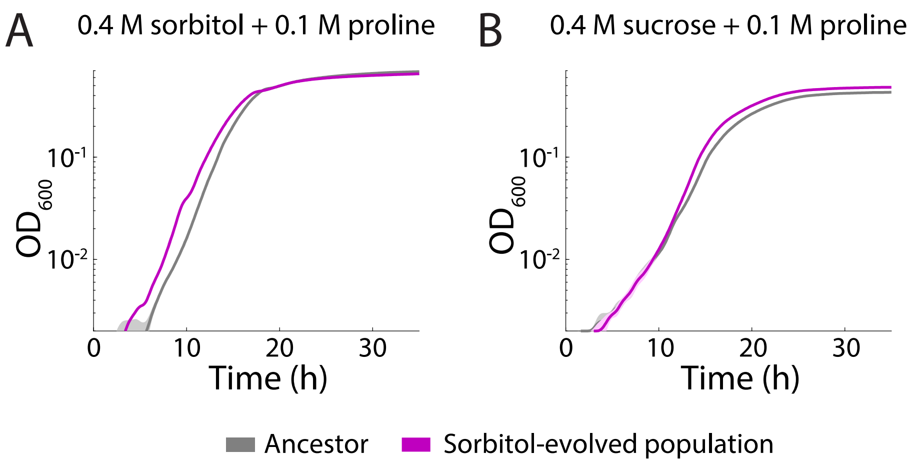


**Supplementary Figure 5: The sorbitol-evolved population still exhibits higher fitness than the ancestor at high-osmolarity when proline is provided as an osmoprotectant.**

A) Growth curves of the ancestor (gray) and So0.5-1 population (magenta) grown in DM25 supplemented with 0.4 M sorbitol and 0.1 M proline. Growth curves are averages of *n*=3 replicates, with shaded regions representing the standard error of the mean (SEM). The SEM is small at some time points and is partially obscured by the line.

B) Growth curves of the ancestor (gray) and So0.5-1 population (purple) grown in DM2500 supplemented with 0.4 M sucrose and 0.1 M proline. Growth curves are averages of *n*=3 replicates, with shaded regions representing the SEM. The SEM is small at some time points and is partially obscured by the line.

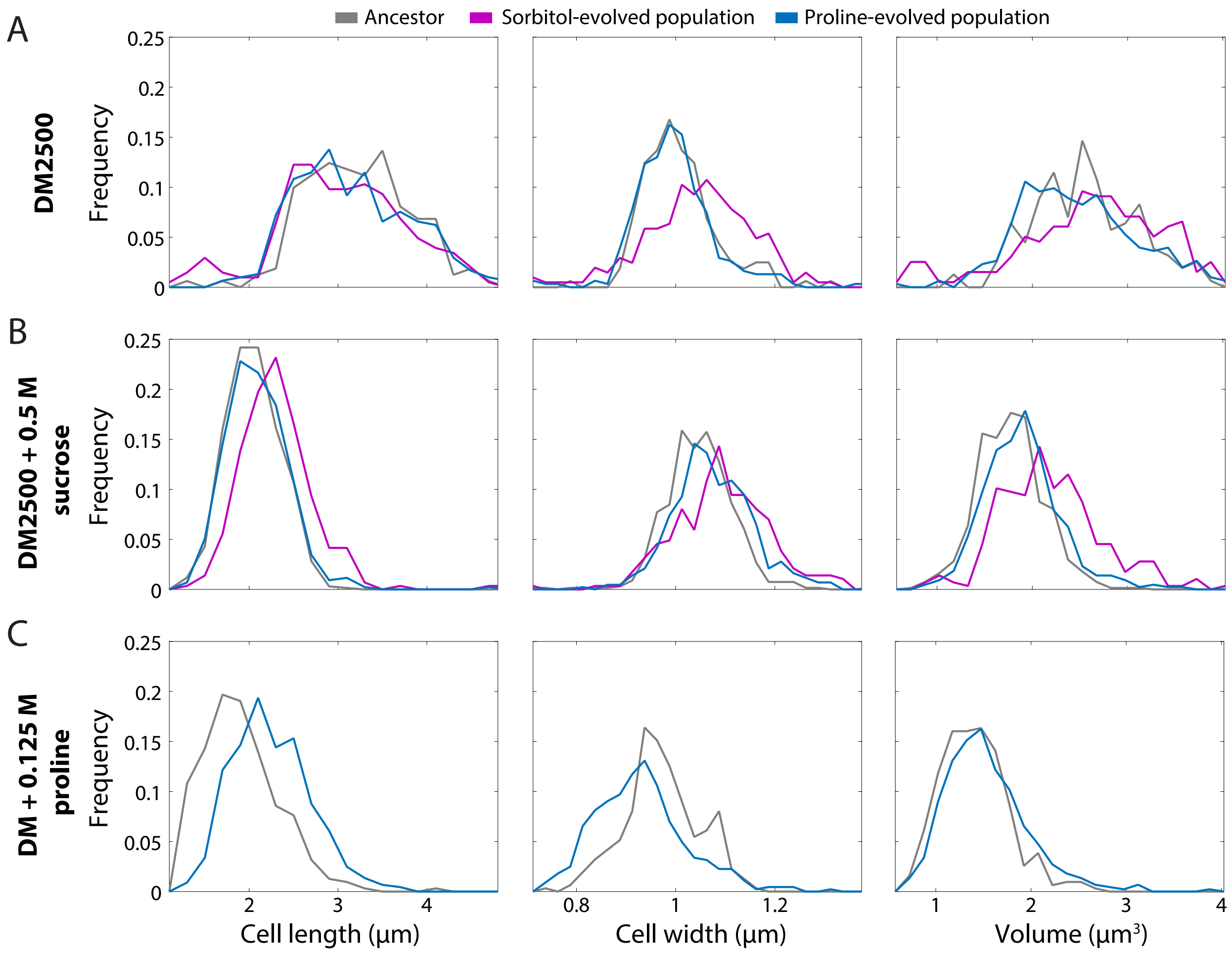


**Supplementary Figure 6: Cell shape distributions of the evolved populations and the ancestor.**

Distributions of cell width, length, and volume in exponential phase were measured for the ancestor, So0.5-1 population, and Pr0.5-1 population grown in DM2500 (A), DM2500+0.5 M sucrose (B), and DM+0.125 M proline (C) (*n*>160 cells for each population).

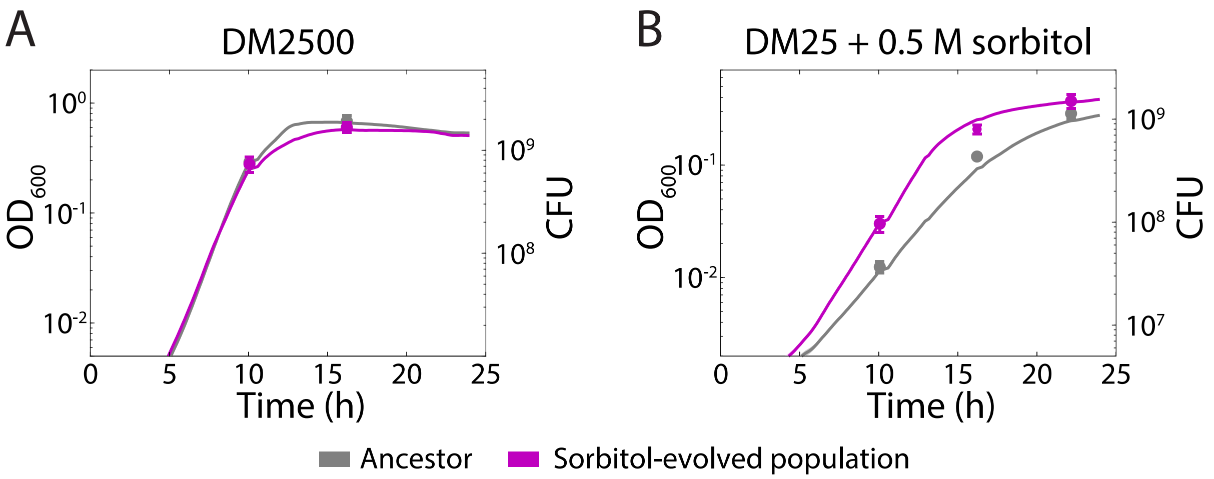


**Supplementary Figure 7: CFU measurements validate OD-based conclusions regarding the relative fitness of the sorbitol-evolved population compared with the ancestor.**

A) OD growth curves (lines) and CFUs (circles and error bars, $n$=3) during growth in DM2500 were highly overlapping for both the ancestor (gray) and So0.5-1 population (magenta). Growth curves are averages of $n$=15 replicates, with shaded regions representing the standard error of the mean (SEM). The SEM is small at many time points and is partially obscured by the line.

B) OD growth curves (lines) and CFUs (circles and error bars, $n$=3) during growth in DM25 supplemented with 0.5 M sorbitol were highly overlapping for both the ancestor (gray) and So0.5-1 population (magenta). Growth curves are averages of $n$=15 replicates, with shaded regions representing the SEM. The SEM is small at many time points and is partially obscured by the line.
